# Transcriptional network involving ERG and AR orchestrates Distal-Less Homeobox 1 mediated prostate cancer progression

**DOI:** 10.1101/2020.08.28.271916

**Authors:** Sakshi Goel, Vipul Bhatia, Shannon Carskadon, Nilesh Gupta, Mohammad Asim, Nallasivam Palanisamy, Bushra Ateeq

## Abstract

Nearly half of the prostate cancer (PCa) cases show elevated levels of ERG oncoprotein due to *TMPRSS2-ERG* gene fusion. Here, we demonstrate ERG mediated upregulation of Distal-less homeobox-1 (DLX1), an established PCa biomarker. Using series of functional assays, we show DLX1 elicits oncogenic properties in prostate epithelial cells, and abrogating its function leads to reduced tumor burden in mouse xenografts. Clinically, ∼60% of the PCa patients exhibit high DLX1 levels, while ∼50% of these cases also harbor elevated ERG associated with aggressive disease and poor survival probability. Mechanistically, we show that ERG gets recruited onto *DLX1* promoter and interacts with its enhancer-bound androgen receptor (AR) and FOXA1 to regulate *DLX1* expression in *TMPRSS2-ERG* positive cases. Alternatively, in ERG-negative cases, DLX1 is regulated by AR/AR-V7 and FOXA1. Importantly, BET bromodomain inhibitors disrupt the transcriptional regulators of *DLX1* and its associated oncogenic properties, signifying their efficacy in treatment of DLX1-positive PCa patients.

## Introduction

Gene rearrangement involving the androgen-regulated transmembrane protease Serine 2 (*TMPRSS2)* and ETS transcription factor, *ERG* (v-ets erythroblastosis virus E26 oncogene homolog) recurrent in ∼50% of the prostate cancer (PCa) patients is often considered as an early event in tumorigenesis, which either alone or in cooperativity with additional genetic alterations such as *PTEN* loss, promotes prostate adenocarcinoma (King et al., 2009; Klezovitch et al., 2008; Tomlins et al., 2005). Aberrant overexpression of ERG oncoprotein controls a transcriptional network linked to PCa development (Chng et al., 2012; Yu et al., 2010), increased metastatic potential, and poor clinical outcome (Mehra et al., 2008; Mosquera et al., 2008). Moreover, ERG is also known to induce EZH2-mediated epigenetic silencing resulting in stem-cell-like dedifferentiation program and inhibits androgen receptor (AR)-mediated gene regulation (Yu et al., 2010). In *TMPRSS2-ERG* positive PCa, ERG binding onto SOX9 promoter opens up cryptic AR binding sites on the SOX9 enhancer thereby regulating its expression, while the loss of SOX9 results in reduced ERG-mediated oncogenicity (Cai et al., 2013). Nonetheless, other critical partners that govern ERG-mediated oncogenesis remain largely unexplored.

Considering the critical role of AR signaling in the development and progression of PCa, androgen deprivation therapy (ADT) wherein inhibitors against enzymes involved in androgen synthesis (abiraterone acetate) or AR antagonists (bicalutamide, enzalutamide) are used as the first-line of treatment for advanced stage PCa patients (Watson, Arora, & Sawyers, 2015). Although, prolonged administration of ADT results in an inevitable cancer relapse due to drug selection pressure, eventually progressing to aggressive castration-resistant prostate cancer (CRPC) stage (Watson et al., 2015). Mounting evidence show sustained AR activity in CRPC, owing to numerous mechanisms including AR amplification, *AR* gene mutations, intra-tumoral androgen synthesis, and expression of constitutively active AR splice variants (AR-Vs) (Shafi, Yen, & Weigel, 2013). Moreover, patients treated with enzalutamide and abiraterone showed increased levels of AR-V7 in circulating tumor cells, and shorter PSA progression–free survival as compared to AR-V7 negative cases (Antonarakis et al., 2014). Unlike full-length AR (AR-FL), the splice variant AR-V7 functions in a ligand-independent manner and remains constitutively active contributing to the androgen-independent growth and disease progression (Hu et al., 2009). Interestingly, AR-V7 is known to heterodimerize with AR-FL and regulate AR target genes in CRPC (D. Xu et al., 2015), while a recent study shows a distinct AR-V7 cistrome which governs cell-context–dependent transcriptomes (Chen et al., 2018).

The *Distal-less homeobox genes (DLX)* belongs to the homeobox containing family of transcription factors (TFs), which are structural homologs of *Drosophila distal-less (Dll). DLX1* being a member of *DLX* family plays an essential role in the development of craniofacial features, jaw and GABAergic (gamma-aminobutyric acid) interneuron (Panganiban & Rubenstein, 2002). Deregulation of various homeobox genes has been linked to several human malignancies including prostate (Abate-Shen, 2002). In hematopoietic cells, DLX1 impedes downstream TGF-β mediated signaling pathways by interacting with Smad4 (Chiba et al., 2003). In PCa, DLX1 is known to functionally interact with β-catenin and regulates downstream β-catenin/TCF4 signaling pathway (Liang et al., 2018). Furthermore, *DLX1* as a biomarker of PCa has been validated across clinically-independent cancer cohorts, wherein *HOXC6* and *DLX1* along with established PCa risk factors accurately predict high-grade disease (Van Neste et al., 2016). Despite DLX1 being a well-established PCa biomarker, the regulatory mechanism involved in its upregulation and its functional significance in the disease progression is poorly understood.

Here, we uncovered transcriptional regulators that govern the expression of DLX1 in PCa. We show that an increased level of DLX1 is positively associated with *ERG* oncoprotein, where ERG positively regulates *DLX1* expression in *ERG* fusion-positive PCa. Additionally, AR along with FOXA1 mediates *DLX1* regulation in ERG-dependent and -independent manner, and hence sensitizes DLX1 expressing PCa cells to bromodomain and extraterminal (BET) domain inhibitor (JQ1). We also identified DLX1-mediated downstream biological processes and pathways that operate in PCa carcinogenesis. Collectively, this study provides mechanistic insight into the DLX1 mediated oncogenesis and reveals the role of ERG, AR and FOXA1 as key transcriptional regulators of *DLX1* in a subset of PCa. Therefore, therapeutic targeting of DLX1 represents an attractive strategy to inhibit the tumorigenesis of both *TMPRSS2-ERG* positive as well as negative PCa.

## Results

### DLX1 imparts oncogenic properties and promotes PCa progression

The utility of *DLX1* as an non-invasive diagnostic biomarker in PCa patients has been well established (Van Neste et al., 2016), however, its role in pathogenesis of this disease remains unexplored. To establish the association between increased DLX1 levels and PCa progression, we analyzed RNA-sequencing (RNA-Seq) data using publicly available The Cancer Genome Atlas Prostate Adenocarcinoma (TCGA-PRAD) dataset. Interestingly, higher expression of *DLX1* was observed in patients with primary PCa as compared to healthy men (*Figure 1A*). Similarly, in other clinical genomics datasets (GSE35988 and GSE80609), an increased expression of *DLX1* transcript was observed with increased cancer aggressiveness as compared to benign cases (*Figure 1B and Figure supplement 1A*), indicating its potential association with cancer progression. In agreement to this, patients with higher DLX1 expression (DLX1^Hi^) experienced poor survival probability compared to those with lower DLX1 expression (DLX1^Lo^) (*Figure 1C*). Importantly, elevated DLX1 levels were observed in PCa cell lines (22RV1, VCaP, and PC3) that represent CRPC as compared to LNCaP, an androgen-responsive, and RWPE1, a benign and immortalized prostate epithelial cell line (*Figure supplement 1B*). Also, the read coverage of *DLX1* transcript from RNA-seq data of RWPE1, 22RV1, and VCaP cell lines showed similar trends (*Figure 1D*). To understand the functional significance of DLX1, we ectopically overexpressed DLX1 in RWPE1 cells and confirmed its overexpression both at transcript and protein levels (*Figure supplement 1C*). Interestingly, a significant increase in cell proliferation was observed in RWPE1-DLX1 cells as compared to control (*Figure 1E*). Likewise, DLX1 overexpression markedly increased foci forming ability and migratory properties of RWPE1 cells (*Figure 1F and 1G*). Contrariwise, to understand the functional role of DLX1 using loss-of-function model, CRISPR/Cas9 mediated gene knockout of *DLX1* was performed in 22RV1 cells (*Figure supplement 1D*), and *DLX1*-knockout was confirmed by genomic amplification of the CRISPR/Cas9 target gene sequence (*Figure supplement 1E*). Three independent 22RV1-*DLX1*-KO clones (C-1, C-2, C-3) showing loss of DLX1 expression both at transcript and protein levels (*Figure supplement 1F*) were examined to determine any phenotypic changes in these genetically engineered lines. Notably, a marked reduction in proliferation rates of 22RV1-*DLX1-*KO cells was observed compared to control (*Figure 1H*). Similarly, a significant reduction (∼60%) in the cell migratory and foci forming properties was observed in 22RV1-*DLX1*-KO cells (*Figure 1I and Figure supplement 1G*). We also observed a ∼4-5-fold attenuation of anchorage-independent growth of 22RV1-*DLX1*-KO cells compared to control (*Figure 1J*).

**Figure 1.**
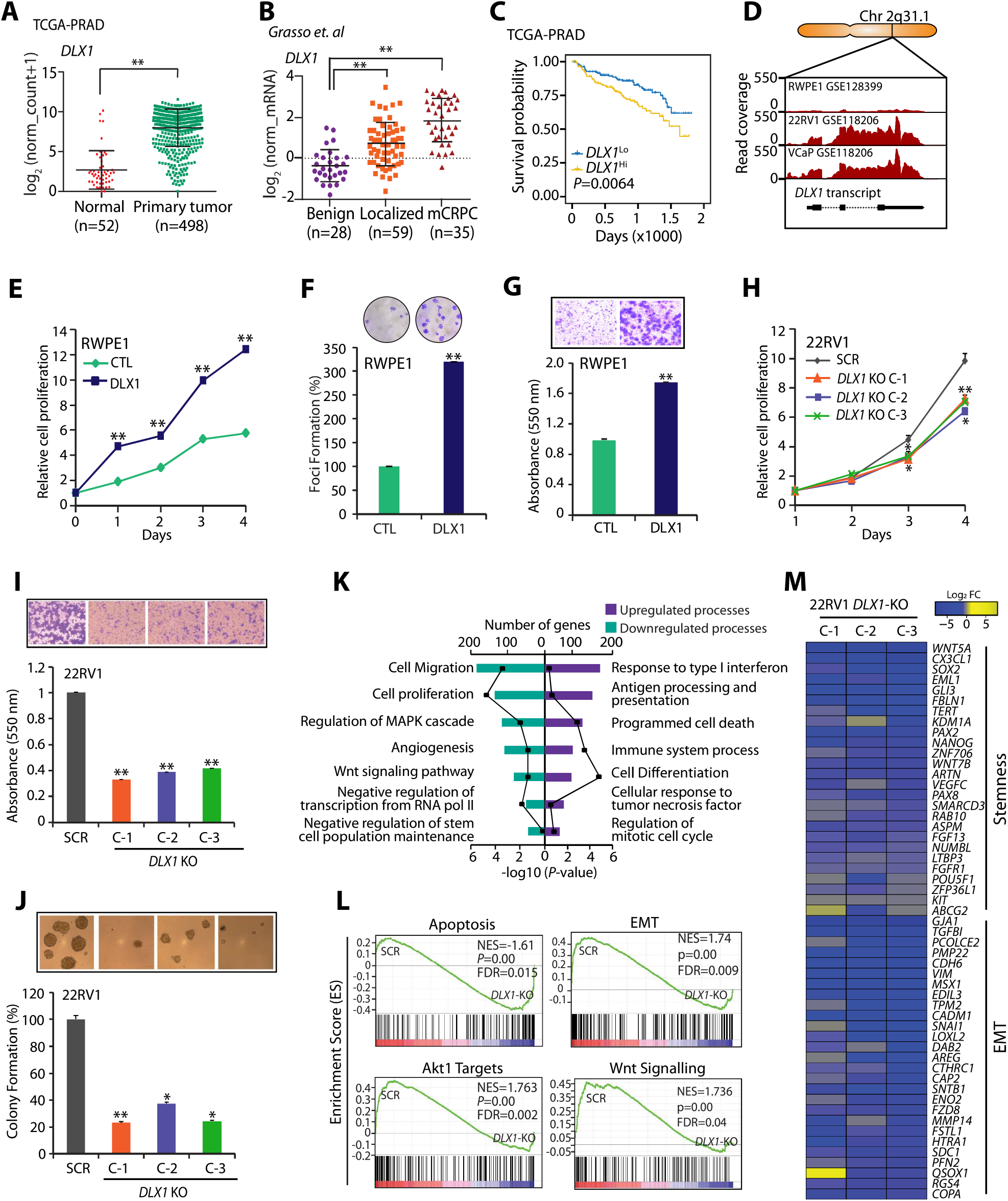
High DLX1 expression associates with poor prostate cancer prognosis and promotes disease progression. **(A)** Dot plot of TCGA-PRAD RNA-seq data showing *DLXl* expression, data represents mean ± SD. Unpaired Student’s t-test was applied. **(B)** Dot plot of DLX1 expression using microarray profiling data (GSE35988), data represents mean ± SD. One-way ANOVA with post hoc Dunnett’s multiple comparisons test was applied. **(C)** Kaplan-Meier plot showing survival probability in TCGA-PRAD (n=498) categorized in high *DLXl (DLX1*^*Hi*^) and low *DLXl (DLX1*^*Lo*^) expression. **(D)** RNA-seq data showing *DLXl* transcript read counts using publicly available datasets (GSE128399 and GSEl 18206). **(E)** Cell proliferation assay using RWPE1-DLX1 overexpressing cells at indicated time-points. **(F)** Foci formation assay using same cells as in (E). Inset shows representative image of foci formed. **(G)** Boyden Chamber Matrigel migration assay using same cells as in (E). Inset shows representative image of migrated cells. **(H)** Cell proliferation assay using 22RV1-DLX7-KO and control cells at indicated time-point s. (I) Boyden Chamber Matrigel migration assay using same cells as in (H). Inset shows representative images of migrated cells. **(J)** Anchorage-independent soft agar assay using same cells as in (H). Inset shows representative images of colonies formed. **(K)** DAVID analysis showing upregulated (right) and downregulated (left) biological processes in 22RV1*-DLXl-KO* cells versus control. Bars represent the −log10 (P-value) and the frequency polygon (black line) signifies number of genes. **(L)** Same as in (K), except GSEA enrichment plot representing deregulated pathways. **(M)** Heatmap displaying downregulated genes involved in cancer sternness and EMT using 22RV1-DLX7-KO cells. In the panels (E-J), data shown from three biologically independent samples (n=3). Data represent mean ± SEM. * *P*≤0.05 and \*\**P*≤ 0.005 using two-tailed unpaired Student’s t-test.

To delineate the biological pathways involved in DLX1-mediated oncogenesis, we performed microarray-based global gene expression profiling of 22RV1-*DLX1*-KO and 22RV1-SCR cells. Database for Annotation, Visualization and Integrated Discovery (DAVID) bioinformatics analysis on the differentially expressed genes revealed important up- and down-regulated biological processes engaged by DLX1 (Supplementary Table S1). As anticipated, genes involved in proliferation and migration of cells, angiogenesis, Wnt signaling, and stem-cell population maintenance were significantly downregulated in 22RV1-*DLX1*-KO cells (*Figure 1K*). Conversely, genes associated with cell cycle regulation, antigen processing and presentation, immune system process and response to tumor necrosis factor and type I interferon were found to be upregulated (*Figure 1K*). Subsequent gene set enrichment analysis (GSEA) showed enrichment of genes involved in the apoptotic pathway in *DLX1*-KO cells, whereas genes involved in epithelial-to-mesenchymal transition (EMT), active Akt and Wnt signaling were enriched in the control 22RV1 cells (*Figure 1L*). Additionally, we observed differentially regulated genes upon *DLX1*-KO were involved in EMT and cancer stemness (*Figure 1M*). Thus, our findings show the significance of DLX1 in imparting oncogenic properties to prostate cells.

### DLX1 elicits biological processes involved in cancer progression and tumorigenesis

To further explore role of DLX1 in regulating oncogenic pathways, we examined expression of EMT genes at transcript and protein level. As anticipated, reduced expression of two archetypal markers associated with mesenchymal phenotype (vimentin and snail), while increased expression in E-cadherin, an epithelial marker in *DLX1*-KO cells was observed (*Figure 2A*). This finding was reinforced by the reduced vimentin levels in 22RV1-*DLX1*-KO cells, contrariwise the increased expression was observed in DLX1 overexpressing RWPE1 cells (*Figure 2B and Figure supplement 1H*). In accordance with the microarray data, we observed a decrease in the expression of established stem cell markers including *POU5F1* (Oct-4), *ABCG2*, *CD117* (c-kit) and *SOX2* (*Figure 2C*). We also examined the cell surface expression of two well-known stem cell markers, ABCG2 and CD44 in *DLX1*-KO cell lines. Interestingly, a marked reduction in the expression of ABCG2 (∼30-60%) and CD44 (∼50-80%) was observed in 22RV1-*DLX1*-KO cells, while a robust increase in the expression of these markers was noted in RWPE1-DLX1 cells (*Figure 2D*). Since increased aldehyde dehydrogenase 1A1 (ALDH1A1) activity is often associated with cancer stem cell phenotype (Li et al., 2010), we performed ALDH activity assay on 22RV1-*DLX1*-KO cells that showed a ∼40% reduction in the ALDH activity (*Figure 2E and Figure supplement 1I*). Conversely, RWPE1-DLX1 cells showed a significant increase in the ALDH activity, suggesting a potential role of DLX1 in promoting cancer stemness (*Figure 2F and Figure supplement 1J*). Next, the role of DLX1 in regulating cell cycle was assessed using flow-cytometry based analysis. A significant increase in percentage of cells arrested in the S-phase with a concomitant decrease in G2/M phase was observed upon *DLX1*-knockout compared to control cells (*Figure 2G and Figure supplement 1K*). Subsequently, an increase in pAkt levels was observed in the RWPE1-DLX1 cells (*Figure supplement 1H*), indicating the critical role of DLX1 in regulating cancer cell survival and proliferation. Consistent with the GSEA data, increased expression of the late apoptotic markers was observed as indicated by an increase in the number of Annexin-V and 7-AAD (7-amino-actinomycin D) stained *DLX1*-KO cells (*Figure 2H*). The increased apoptosis observed in *DLX1*-KO cells was associated with Caspase 3 activation induced Poly-(ADP-ribose) polymerase (PARP) cleavage that can trigger apoptosis. Also, decrease in the levels of anti-apoptotic Bcl-xL protein was observed (*Figure 2I*). Differentially regulated genes in the *DLX1*-KO clones were functionally characterized by forming an overlapping network using Enrichment map that revealed EMT, cell cycle and DNA damage as the major downregulated pathways (*Figure supplement 2*). Further to validate the oncogenic role of DLX1 in another aggressive PCa cells, we analyzed the publicly available RNA-seq data for transient knockdown of *DLX1* in LNCaP derivative osteotropic cell line C4-2B (GSE78913) (Rhie et al., 2016). As anticipated DLX1 was one of the top 5 downregulated genes in the volcano plot depicting differentially regulated genes (*Figure 2J*). Pathway analysis of differentially regulated genes showed significant downregulation of the genes involved in cell division, NF-κB signalling and MAP kinase cascade while genes associated with apoptosis, cell-cell adhesion and cell-cycle arrest were upregulated (*Figure 2K*). Additionally, GSEA analysis of the deregulated genes showed significant decrease in androgen response and E2F pathway consequent to *DLX1* silencing in C4-2B cells (*Figure 2L*). Thus, reaffirming the role of DLX1 in eliciting oncogenic phenotype in PCa cells.

**Figure 2.**
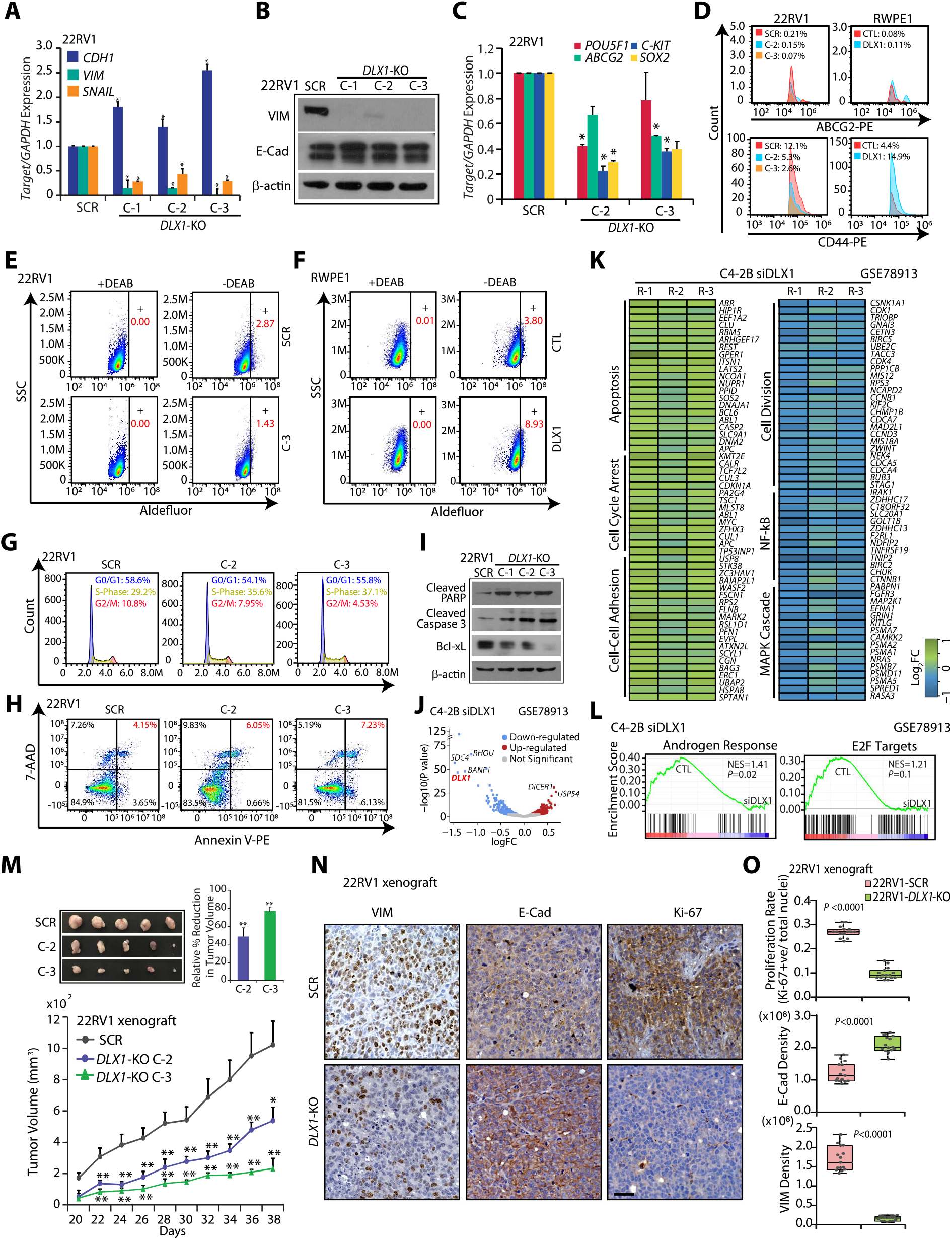
Ablating DLX1 expression inhibits oncogenic properties and tumorigenesis in mice model. **(A)** Q-PCR for EMT markers in 22RV1-OLX7-KO and control cells. **(B)** lmmunoblot for vimentin (VIM) and E-Cad using same cells as in (A). 1,-actin was used as positive control. **(C)** Q-PCR analysis for stem cell marker genes in *DLX1-KO* and control cells. **(D)** Histogram showing flow cytometry data depicting ABCG2 (top) and CD44 (bottom) expression in 22RV1-OLX7-KO (left), and RWPEl-DLXl overexpressing (right) cells. **(E)** Fluorescence intensity of catalyzed ALDH substrate in *DLX1-KO* and control cells. Marked windows show ALDHl + percent cell population **(F)** Same as in (E) except for RWPEl-DLXl overexpressing and control cells. **(G)** Flow cytometry data for cell cycle distribution of 22RV1-OLX7-KO and control cells. **(H)** Flow cytometry-based apoptosis assay using 22RV1-OLX7-KO cells stained with Annexin V-PE and 7-AAD. **(I)** lmmunoblot showing cleaved PARP, cleaved Caspase-3 and Bcl-xl **(J)** Volcano plot showing deregulated genes upon *siDLX1* in C4-2B cells in publicly available RNA-seq dataset (GSE78913). **(K)** Heatmap displaying deregulated genes involved in biological processes using same cells as in (J). R-1, R-2 and R-3 represents three independent replicates. **(L)** Same as in (J) except GSEA plots showing deregulated path ways. **(M)** Tumor volume of 22RV1-OLX7-KO and control cells subcutaneously implanted in NOD/SCIO mice (n=6). Representative images of the tumors harvested at end of the experiment are shown as inset (left). Bar graph showing relative percent reduction in tumor burden showed in inset (right). **(N)** Representative images depicting tumor xenografts stained for Ki-67, E-Cad, and VIM by IHC. Scale bar, 35μm. **(0)** Ki-67, E-Cad and VIM positive cells were quantified from 15 random histological fields. Data shown from three biologically independent sa mples. In the panels (A), (C-G), and (1) data represent mean ± SEM. \**P*≤ 0.05 and ** *P*≤ 0.005 using two-tailed unpaired Student’s t-test.

Further, to establish the role of DLX1 in tumorigenesis, we subcutaneously implanted 22RV1-*DLX1*-KO and 22RV1-SCR control cells into the flank region of immunodeficient NOD/SCID mice and monitored the tumor growth. Remarkably, a significant reduction (∼80%) in tumor burden was observed in the mice implanted with 22RV1-*DLX1*-KO cells as compared to the control group (*Figure 2M*). Reduced expression of Ki-67 further confirms decrease in cell proliferation rate of 22RV1-*DLX1*-KO cells in xenografted mice. Also, increase in E-Cadherin and reduction in Vimentin expression was observed in mice xenograft comprising *DLX1*-KO cells compared to control group (*Figure 2N and 2O*). Taken together, we demonstrate that increased DLX1 expression confers EMT, cancer stemness, and promotes tumor progression.

### High level of ERG oncoprotein shows positive association with DLX1 expression

Overexpression of the ERG TF due to *TMPRSS2–ERG* genetic rearrangement is considered an early event in disease pathogenesis in about ∼50% of PCa cases (Tomlins et al., 2005), thus we next interrogated association of DLX1 with ERG using publicly available TCGA-PRAD cohort. We stratified clinical genomic data of patients (n=498) based on the expression of *ERG,* and intriguingly most of the cases with higher *ERG* expression exhibits increased expression of *DLX1* transcript (*Figure 3A*). Next, using the UALCAN cancer OMICS database (Chandrashekar et al., 2017) we analyzed the overall survival probability of patients with varying expression of *DLX1* and its association with the well-known seven molecular PCa subtypes defined by TCGA (Abeshouse et al., 2015). Notably, *TMPRSS2-ERG* fusion-positive patients with higher levels of DLX1 experienced lower survival probability as compared to fusion-positive patients with low/medium *DLX1* expression (*Figure supplement 3A*), indicating the existence of oncogenic cooperativity between DLX1 and ERG oncoprotein. Furthermore, a significant positive correlation between *DLX1* and *ERG* expression was observed in both MSKCC (Taylor et al., 2010) and TCGA-PRAD cohorts (*Figure 3B and Figure supplement 3B*).

**Figure 3.**
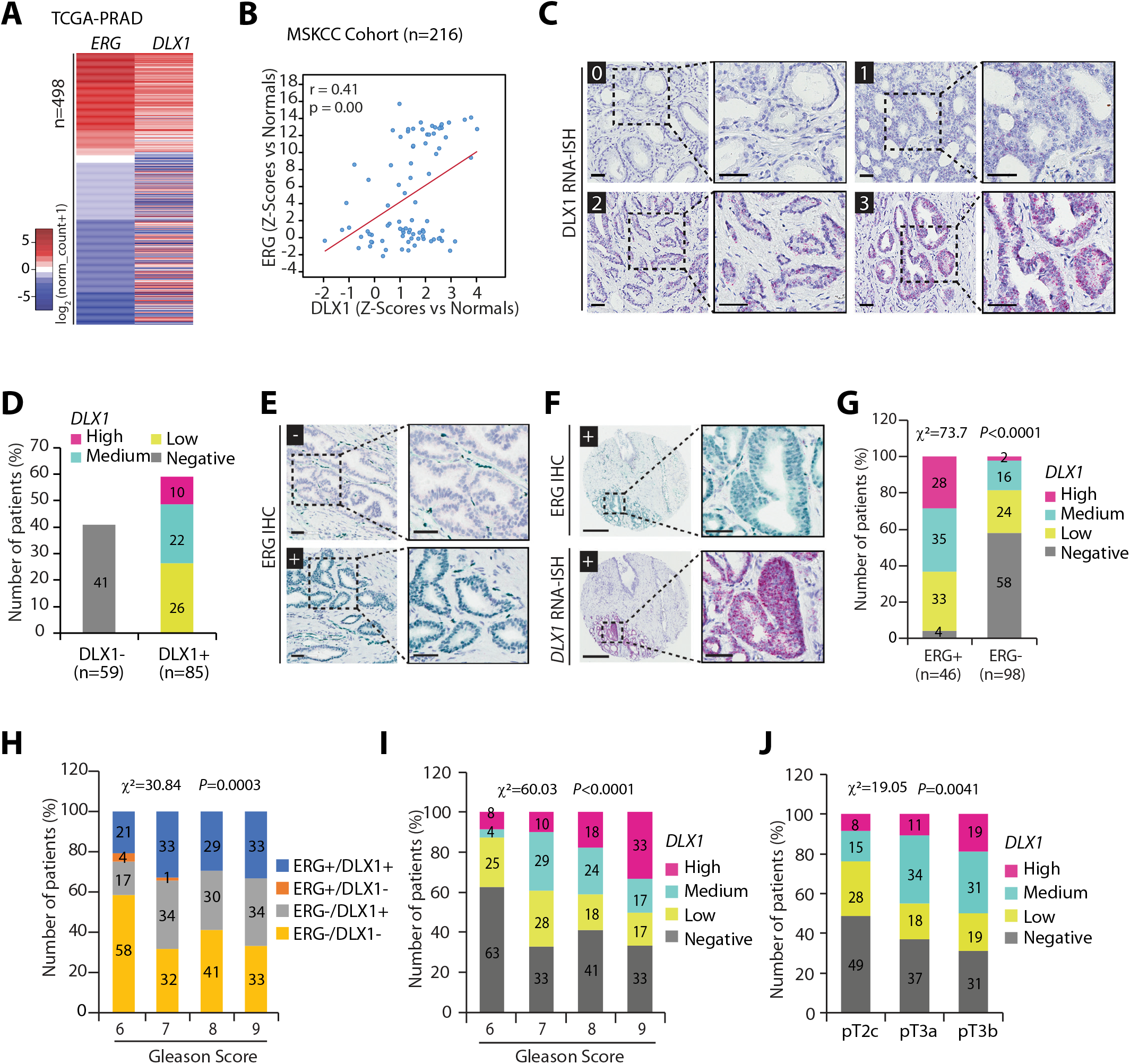
ERG-positive prostate cancer patients associate with increased DLX1 levels and aggressive disease. **(A)** Heatmap generated using TCGA-PRAD RNA-seq data for *ERG* and *DLX1* in primary tumor PCa cases (n=498). Shades of red and blue represent the log2 (norm_count+1) value. **(B)** Correlation plot between *DLX1* and *ERG* using Memorial Sloan Kettering Cancer Center cohort (MCKCC) dataset available at cBioPortal. **(C)** Representative foci of PCa TMA cores (n=144) showing RNA-ISH scoring pattern of *DLX1*, score 0 represents *DLX1* negative, score 1 signifies low *DLX1*, score 2 indicates medium *DLX1* and score 3 represents the high *DLX1* expression. Scale bar, 50μm. **(D)** Bar plot showing percent patients negative (DLX1-) and positive (DLX1+) for DLX1 expression based on the scoring pattern. **(E)** Same as in (C) except IHC staining for ERG. Scale bar, 50μm. **(F)** Same as in (C) except IHC for ERG (top) and RNA-ISH for *DLX1* (bottom). Scale bar for (E-F) is represented as 300μm and 50μm for the entire core and inset image, respectively. **(G)** Bar plot showing percent patients with varying *DLX1* expression in ERG-positive (ERG+) and -negative (ERG-) PCa cases. **(H)** Same as in (G) except for patients showing association of ERG and *DLX1* expression status to patients’ Gleason scores. **(I)** Same as in (G), except association between *DLX1* expression and patients’ Gleason scores. **(J)** Same as in (G), except association of *DLX1* with tumor stage. For panels (G-J) *P*-value were calculated using Chi-Square test.

To further support these findings, we performed IHC, and RNA-ISH for ERG and *DLX1* expression, respectively using TMA comprising 144 PCa patients. Except three cases, none of these patients were administered with hormone therapy. *DLX1* RNA-ISH staining patterns were classified into four levels ranging from the score of 0 to 3, and nearly 60% of patients were positive for *DLX1* expression ranging from low to high (*Figure 3C and 3D*). Specimens stained for ERG were stratified into ERG positive (ERG+) and negative (ERG-) categories (*Figure 3E*). In concordance with our *in-silico* analysis, 44 out of 46 *TMPRSS2-ERG* fusion-positive cases (∼96%) showed positive staining for *DLX1* expression (*Figure 3F, 3G, and Figure supplement 3C*), wherein ∼28% patients showed a high *DLX1* (*DLX1*^Hi^, score 3), ∼35% patients with moderate (*DLX1*^Me^, score 2) and ∼33% patients with lower *DLX1* expression (*DLX1*^Lo^, score 1) (*Figure 3G*). In contrast, among ERG-negative tumors only ∼42% were positive for *DLX1* expression, suggesting a role for ERG TF in regulating *DLX1* (*Figure supplement 3C*). In terms of clinical staging, the percentage of tumors stained positive for both ERG and *DLX1* increased from 21% in low Gleason score (GS6) to 33% in high Gleason score disease (GS9) (*Figure 3H*). Moreover, DLX1 expression alone gradually increased as a function of disease stage; 37% positive in GS6 disease to 67% positive in GS9 disease (*Figure 3I*), similarly, 51% in pT2c (pathologic tumor 2c) to 69% in pT3b stage (*Figure 3J*). Thus, suggesting that the majority of the patients harboring higher *DLX1* and/or ERG expression (ERG+/DLX1+ and ERG-/DLX1+) were associated with advanced PCa. Together this data suggests the role of DLX1 in aggressive tumor stage and accentuate the possible oncogenic cooperativity between DLX1 and ERG in the PCa progression.

### ERG transcriptionally regulates *DLX1* in *TMPRSS2-ERG* positive prostate cancer

Given the strong association between ERG and DLX1 expression in PCa patients, we next investigated the role of ERG in regulating *DLX1* expression. We analyzed publicly available chromatin immunoprecipitation sequencing (ChIP-seq) datasets for ERG in VCaP (*TMPRSS2-ERG* fusion-positive) cell line, and an increased enrichment of ERG on the *DLX1* promoter was observed, while ERG occupancy was reduced in small hairpin RNA (shRNA) mediated *ERG* depleted cells (*Figure 4A*). Furthermore, enrichment of ERG on the *DLX1* promoter was also observed in ectopically ERG overexpressing RWPE1 cells (*Figure 4A*). Next, we scanned ∼1 kb upstream and 500bp downstream region to the transcription start site (TSS) of *DLX1* for any potential ERG binding motif (EBM), and two putative ERG binding sites namely, EBM1 (P1) and EBM2 (P2) were observed (*Figure 4B*). As expected, our ChIP-qPCR data confirmed the significant occupancy of ERG at these EBMs on *DLX1* promoter in VCaP cells (*Figure 4B*). Importantly, these EBMs were associated with transcriptionally active chromatin as a marked enrichment of RNA-polymerase II (RNA-Pol II) and histone H3 lysine 9 acetylation (H3K9Ac) were observed at these sites (*Figure 4B*). ERG target gene, *PLAU* was used as a positive control. In agreement with this, ectopic overexpression of ERG in benign prostate epithelial RWPE1 cells (RWPE1-ERG) results in a ∼10-fold upregulation of DLX1 both at mRNA and protein levels (*Figure 4C*). Furthermore, RWPE1-ERG and control cells were used to examine the ERG-mediated activation of the *DLX1* promoter by performing luciferase-based reporter assay. As expected, a significant increase in the reporter activity was observed in RWPE1-ERG cells containing a wild-type *DLX1* promoter, while no significant change in the luciferase activity was observed with EBMs mutated *DLX1* promoter (*Figure 4D*).

**Figure 4.**
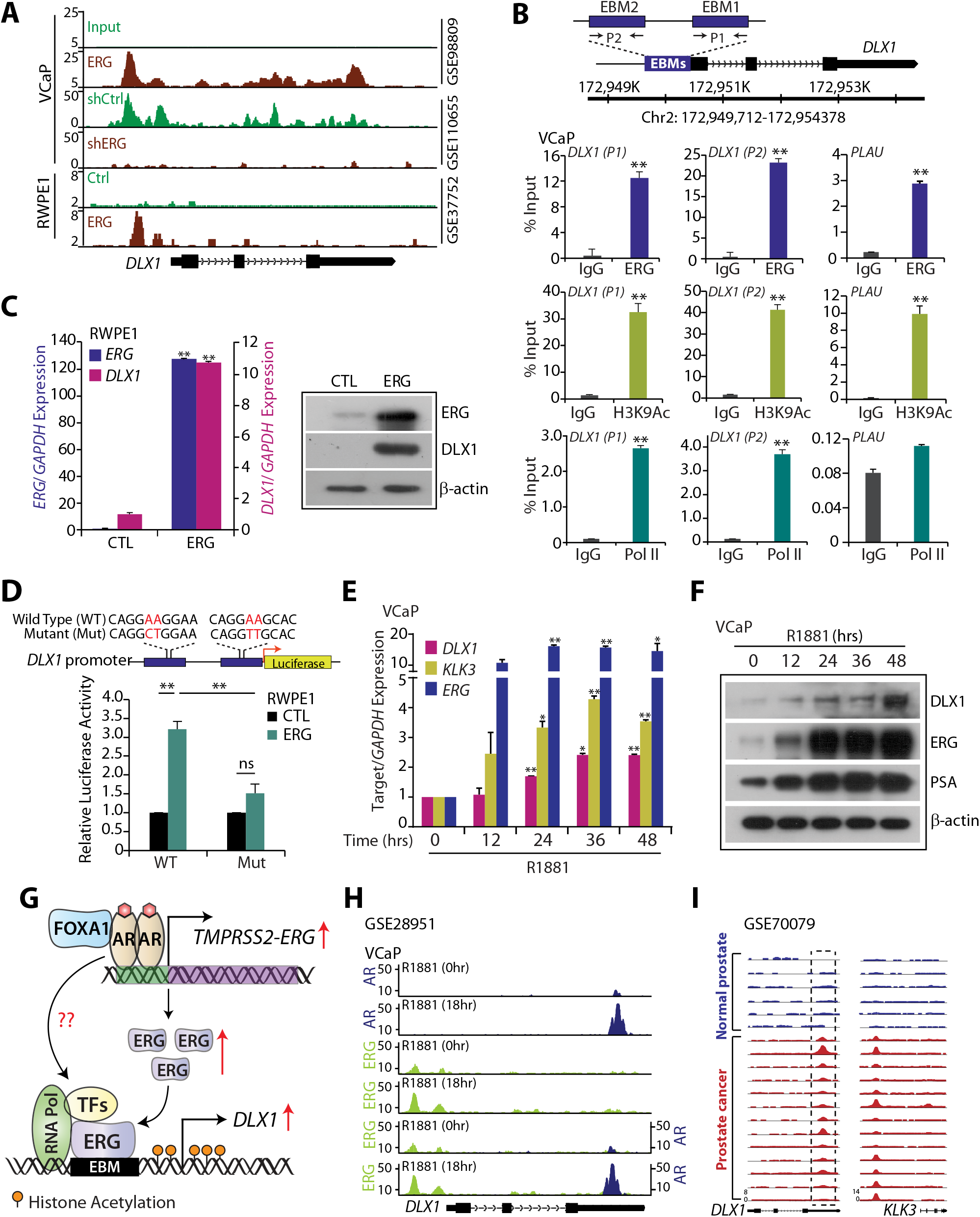
DLX1 is a transcriptional target of ERG in TMPRSS2-ERG positive PCa. **(A)** ChIP-seq data depicting ERG enrichment at *DLX1* promoter in the indicated cell lines using publicly available data from GEO databases (GSE98809, GSE110655 and GSE37752). **(B)** Schema showing chromosomal location of ERG binding motif (EBM) onto DLX1 promoter selected for ChIP-qPCR (top). ChIP-qPCR data showing recruitment of ERG, histone H3 lysine 9 acetylation (H3K9Ac) and RNA-polymerase II (RNA-Pol II) at the *DLX1* and *PLAU* promoter. (**C)** Q-PCR (left) and immunoblot (right) data showing expression of ERG and DLX1 in RWPE1-ERG overexpressing cells. **(D)** Schema showing site-directed mutagenesis of *DLX1* promoter cloned upstream of luciferase gene (top). Luciferase reporter assay in RWPE1 cells demonstrating wild-type (WT) and mutant (Mut) *DLX1* promoter-driven reporter activity in the presence or absence of ERG (bottom). **(E)** Quantitative PCR data showing relative expression of target genes in VCaP cells stimulated with 10nM R1881 at indicated time points. **(F)** Same as (E) except immunoblot data. **(G)** Schematic diagram depicting ERG-mediated transcriptional regulation of *DLX1* and probable role of AR in DLX1 regulation. **(H)** ChIP-seq data showing AR and ERG recruitment in R1881 stimulated VCaP cells (GSE28951). **(I)** ChIP-seq data showing AR enrichment at *DLX1* putative enhancer in the normal prostate (n=6) and PCa (n=13) tissue samples (GSE70079). *KLK3* represents positive control. Data shown from three biologically independent samples. In the panels (B-E), data represent mean ± SEM. \**P*≤ 0.05 and \*\**P*≤ 0.005 using two-tailed unpaired Student’s t-test.

Since AR signaling drives the expression of ERG in *TMPRSS2-ERG* fusion*-*positive background, we next examined the effect of synthetic androgen methyltrienolone (R1881) on the expression of *DLX1* in VCaP cells. Notably, ∼2-fold increase in the DLX1 expression both at mRNA and protein level was observed in R1881-treated VCaP cells (*Figure 4E and 4F*). PSA was used as a positive control for androgen stimulation. Next, we sought to investigate if AR also play a direct role in regulation of *DLX1* transcript (*Figure 4G*). Thus, to ascertain this, we analyzed publicly available ChIP-seq datasets for AR and ERG in R1881 stimulated VCaP cells. Notably, strong binding of AR on the third exon (chr2:172,661,000-172,662,500) of *DLX1* was observed in R1881 stimulated VCaP cells (*Figure 4H*). On the similar lines several studies revealed the presence of exonic enhancers (eExons) can act as a regulatory element for the nearby genes or the host gene on which they resides (Birnbaum et al., 2012; Neznanov, Umezawa, & Oshima, 1997). Hence, we speculate that the presence of AREs at the third exon may act as a putative enhancer region in the transcriptional regulation of *DLX1*. We further examined AR binding on *DLX1* gene using publicly available dataset (GSE70079) comprising normal and PCa specimens, and as expected a remarkable enrichment of AR was observed on the putative enhancer element of *DLX1* in PCa samples (*Figure 4I*). Conclusively, we show that ERG directly gets recruited on the *DLX1* promoter regulating its expression in *TMPRSS2-ERG* positive cases (*Figure 4G*). Also, our findings imply the potential role of AR signaling in mediating *DLX1* expression in prostate cancer.

### Androgen receptor regulates DLX1 expression in prostate cancer

Intrigued by the enrichment of AR at DLX1 enhancer region, we pursued to identify probable association between AR signaling in DLX1 regulation. It has been shown previously that AR binding sites (ARBS) are reprogrammed in tumorous tissues and binds to distinct set of genes compared to normal prostatic tissue (Pomerantz et al., 2015). Notably, DLX1 was found to be in the top 10 differentially upregulated genes in TCGA-PRAD dataset, which harbor tumor-specific ARBS neighboring the 50kb regions (*Figure supplement 4A*). To further validate the AR binding on the putative enhancer of *DLX1*, we performed ChIP-qPCR for AR in R1881-stimulated VCaP cells, and a significant enrichment of AR on *DLX1* gene was observed, which was disrupted upon anti-androgen enzalutamide treatment (*Figure 5A*). The promoter of *KLK3* was used as a positive control (*Figure supplement 4B*). Since, FOXA1 is a known pioneering TF and a coactivator of AR (Jin, Zhao, Wu, Kim, & Yu, 2014), we investigated the occupancy of FOXA1 at the *DLX1* putative enhancer region. Importantly, ChIP-seq analysis of publicly available dataset (GSE56086) reveals FOXA1 binding at the same locus (chr2:172,661,000-172,662,500) as occupied by AR at the putative enhancer of *DLX1* (*Figure supplement 4C*). In corroboration with the above finding ChIP-qPCR in VCaP cells also showed enrichment of FOXA1 on the same position signifying the recruitment of AR transcriptional complex on the *DLX1* putative enhancer region (*Figure supplement 4D*). An AR antagonist EPI-001 is known to inhibit AR activity by targeting its N-terminal domain thereby abrogating AR-V7 mediated transcriptional activity (Myung et al., 2013). In agreement, anti-androgen treatment using enzalutamide and EPI-001 abrogated R1881-induced DLX1 expression in VCaP cells, implicating the role of AR in transcriptional regulation of *DLX1* (*Figure 5B*). Since AR is known to interact with ERG as a coregulatory TF and regulates the expression of target genes (Yu et al., 2010; Zhang et al., 2019), we examined the probable co-interaction of ERG and AR at the promoter of *DLX1* by performing re-ChIP experiment (*Figure 5C*). As anticipated, ChIP using ERG antibody followed by antibody against AR showed significant enrichment of AR and ERG at *DLX1* confirming their interaction via promoter-enhancer region at *DLX1* gene (*Figure 5C*). *CUTL2* was used as a positive control for AR-ERG co-regulated gene. To further validate the probable chromatin interaction at *DLX1* genomic region, we analyzed the 3D-chromatin landscape of RNA-Pol II in VCaP cells using publicly available ChIA-PET dataset (GSE121020) (Ramanand et al., 2020). Also, we performed the integrative analysis of Pol II associated peaks along with ChIP-seq data of DLX1 regulating TFs in PCa. Consistent with our findings, the RNA-Pol II ChIA-PET data confirmed the promoter-enhancer interaction at *DLX1* gene, which also indicated the binding of ERG, AR and FOXA1 transcription factors (*Figure 5D*). Notably, small interfering RNA (siRNA) mediated knockdown of these regulatory factors namely, ERG, AR and FOXA1 in VCaP cells (*Figure supplement 4E*) resulted in reduced DLX1 expression in all the respective conditions. While, more pronounced decrease in the level of DLX1 expression was achieved by simultaneously silencing all three key regulators (*ERG*, *AR* and *FOXA1*), thereby validating the transcriptional interplay between these factors (*Figure 5E and 5F*). Collectively, our results suggest the ERG and AR mediated transcriptional co-regulation of *DLX1* in *TMPRSS2-ERG* fusion-positive cells. Furthermore, shRNA mediated knockdown of *AR* in castration-resistant and *TMPRSS2-ERG* fusion-negative C4-2 cells resulted in decreased DLX1 expression (*Figure supplement 4F*). Moreover, upon androgen stimulation, enrichment of AR and FOXA1 was observed in *TMPRSS2-ERG* fusion-negative 22RV1 cells suggesting the significance of AR in regulating *DLX1* expression in ERG-independent manner. *KLK3* promoter was used as a positive control (*Figure 5G*).

**Figure 5.**
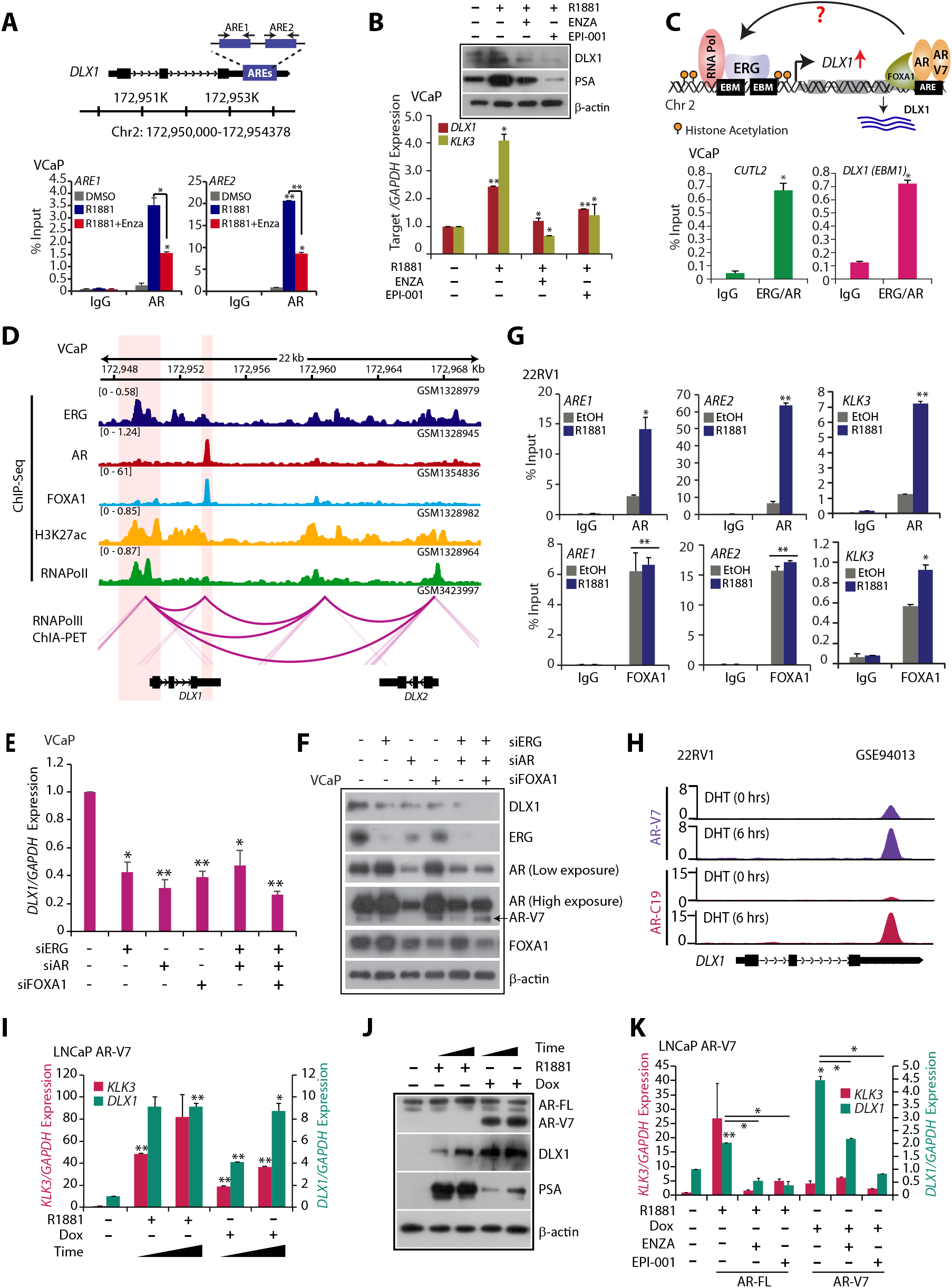
AR and AR-V7 regulate *DLX1* expression in both ERG-dependent and -independent manner. **(A)** Schematic showing the androgen response elements (AREs) at *DLX1* putative enhancer (top). ChIP-qPCR data depicting AR recruitment at *DLX1* putative enhancer in R1881 (10nM) stimulated VCaP cells treated with or without Enzalutamide (10μM). **(B)** Q-PCR and immunoblot showing relative expression of target genes in VCaP cells under culture conditions as mentioned. **(C)** Schematic representation showing the possible interaction between ERG and AR on *DLX1* promoter (Top). Re-ChIP data showing co-enrichment of AR and ERG on EBM at the *DLX1* promoter (Bottom). **(D)** Integrated genome view of 3D chromatin structure and binding of transcription factors at *DLX1* genomic and nearby region. **(E)** Quantitative PCR showing relative expression of *DLX1* in siRNA mediated *AR, ERG* and/or *FOXA1* silenced VCaP cells. **(F)** Same as (E) except for immunoblot data. **(G)** ChIP-qPCR data depicting AR (top) and FOXA1 (bottom) enrichment at *DLX1* putative enhancer in 22RV1 cells stimulated with R1881 (10nM) for 16 hours. *KLK3* is used as a positive control. **(H)** ChIP-seq data (GSE94013) using AR-V7 and AR C-terminal (AR-C19) antibody in DHT stimulated 22RV1 cells. **(I)** Relative expression of *KLK3* and *DLX1* in dox inducible LNCaP AR-V7 overexpressing cells stimulated with R1881 (10nM) and treated with Dox (40ng) or vehicle control for 24 and 48 hours. **(J)** Immunoblot showing expression of AR-FL, AR-V7, DLX1 and PSA using same cells as in (I). β-actin used as loading control. **(K)** Q-PCR showing *DLX1* and *KLK3* expression in LNCaP AR-V7 cells under culture conditions as mentioned for 48 hours. Data shown from three biologically independent samples. In the panels (A-C), (E), (G), (I) and (K), data represent mean ± SEM. *P≤ 0.05 and **P≤ 0.005 using two-tailed unpaired Student’s t-test.

Having established the upregulation of DLX1 in CRPC cases and a substantial role of full-length AR in regulating its expression, we conjecture the possible involvement of AR-V7 an AR splice variant in the upregulation of DLX1 in CRPC cases. Intriguingly, ChIP-seq analysis for AR-V7 specific binding in 22RV1 cells revealed remarkable enrichment of AR-V7 at the putative enhancer of *DLX1* (*Figure 5H*). Moreover, the analysis of publicly available RNA-seq data (GSE94013) shows a significant reduction in *DLX1* transcript upon knockdown of AR-V7 or AR-FL in 22RV1 cells (*Figure supplement 4G*) suggesting the role of AR-V7 in *DLX1* regulation. Furthermore, using an inducible expression system for AR-V7, we generated LNCaP cells that can express both AR-FL and AR-V7, a system that mimics the clinical state in the majority of PCa patients with resistance to enzalutamide/abiraterone (He et al., 2018). In these modified LNCaP cells, the expression levels of AR-FL and AR-V7 can be induced by treating them with R1881 and doxycycline (dox), respectively (*Figure 5I*). In line with this, both AR-FL, as well as AR-V7 expression, led to increased expression of DLX1 at the mRNA and protein levels (*Figure 5I and 5J*). Interestingly, we observed that dox-inducible overexpression of AR-V7 led to much higher expression of DLX1 than by R1881 stimulation alone, suggesting consecutive activation of androgen signaling via AR-V7 in regulating DLX1 expression. In accordance with our hypothesis, treating LNCaP AR-V7 cells with AR antagonists (enzalutamide and EPI-001) in the presence of R1881 or dox abrogated AR-FL and AR-V7 mediated increase in DLX1 expression (*Figure 5K*). Taken together, our findings suggest a role for AR-FL and AR-V7 in the transcriptional regulation of *DLX1* in both ERG-dependent and -independent manner. These findings also highlight the critical role of AR-V7 in the transcriptional regulation of *DLX1*, thereby resonating with relatively high DLX1 expression in PCa patients with advanced stage disease and its association with higher Gleason score.

### BET domain inhibitor attenuates DLX1 expression in prostate cancer

Having established ERG-mediated transcriptional regulation of *DLX1* in *TMPRSS2-ERG* fusion-positive PCa, and significance of AR signaling in regulating DLX1, we next explored the therapeutic strategies to target this regulatory circuitry. Since the utility of BETi, namely JQ1 and I-BET762 has been shown to abrogate aberrant AR signaling and BRD4 localization to AR target genes including *TMPRSS2-ERG* fusion (Asangani et al., 2014). We sought to investigate the efficacy of JQ1 in inhibiting transcriptional regulators of *DLX1*, namely ERG and AR. Using publicly available ChIP-seq dataset (GSE55064), we examined change in ERG recruitment on the *DLX1* promoter in JQ1 treated VCaP cells. Interestingly, a remarkable decrease in the enrichment of ERG was observed in JQ1 treated VCaP cells compared to control. Moreover, the presence of H3K27Ac marks in the control VCaP cells indicate active *DLX1* promoter, and *PLAU*, an ERG target gene was used as a positive control (*Figure 6A*). Further, these results were validated by performing ChIP-qPCR for ERG in JQ1 treated VCaP cells and similar trend was observed (*Figure 6B*). We next investigated change in DLX1 expression in VCaP cells treated with 0.5μM and 1μM of JQ1, and a significant decrease in its expression was observed (*Figure 6C and 6D*). To examine whether JQ1 could inhibit ERG-mediated DLX1 expression, stable RWPE1-ERG cells were treated with JQ1 and a significant decrease in DLX1 expression was observed, thereby confirming the efficacy of JQ1 in inhibiting ERG-mediated transcriptional regulation of *DLX1* (*Figure 6E*). Considering ERG-independent role of AR in *DLX1* regulation, we next treated C4-2, C4-2B cells, and 22RV1 cells with JQ1, and as anticipated a significant reduction in the *DLX1* expression was noted (*Figure 6F*). Furthermore, VCaP and C4-2 cells treated with JQ1 showed a significant decrease in cell proliferation as compared to vehicle control (*Figure 6G*), indicating that JQ1-mediated downregulation of DLX1 might contribute to diminution of oncogenic properties. Taken together, our findings demonstrate ERG and AR-mediated novel regulatory mechanisms responsible for DLX1 upregulation in an aggressive subset of PCa patients. We establish that in *TMPRSS2-ERG* positive PCa, AR along with FOXA1 regulates expression of *DLX1* via interacting with ERG as a coregulatory TF. On other hand, in fusion-negative PCa cases, AR and FOXA1 acts in an ERG-independent manner possibly in association with other coregulatory factors to regulate the expression of DLX1, subsequently imparting DLX1-mediated oncogenesis and disease progression. Importantly, we also demonstrate pharmacological inhibition of ERG/AR-mediated transcriptional circuitry by BET inhibitor results in decrease in DLX1 expression and its oncogenesis (*Figure 6H*).

**Figure 6.**
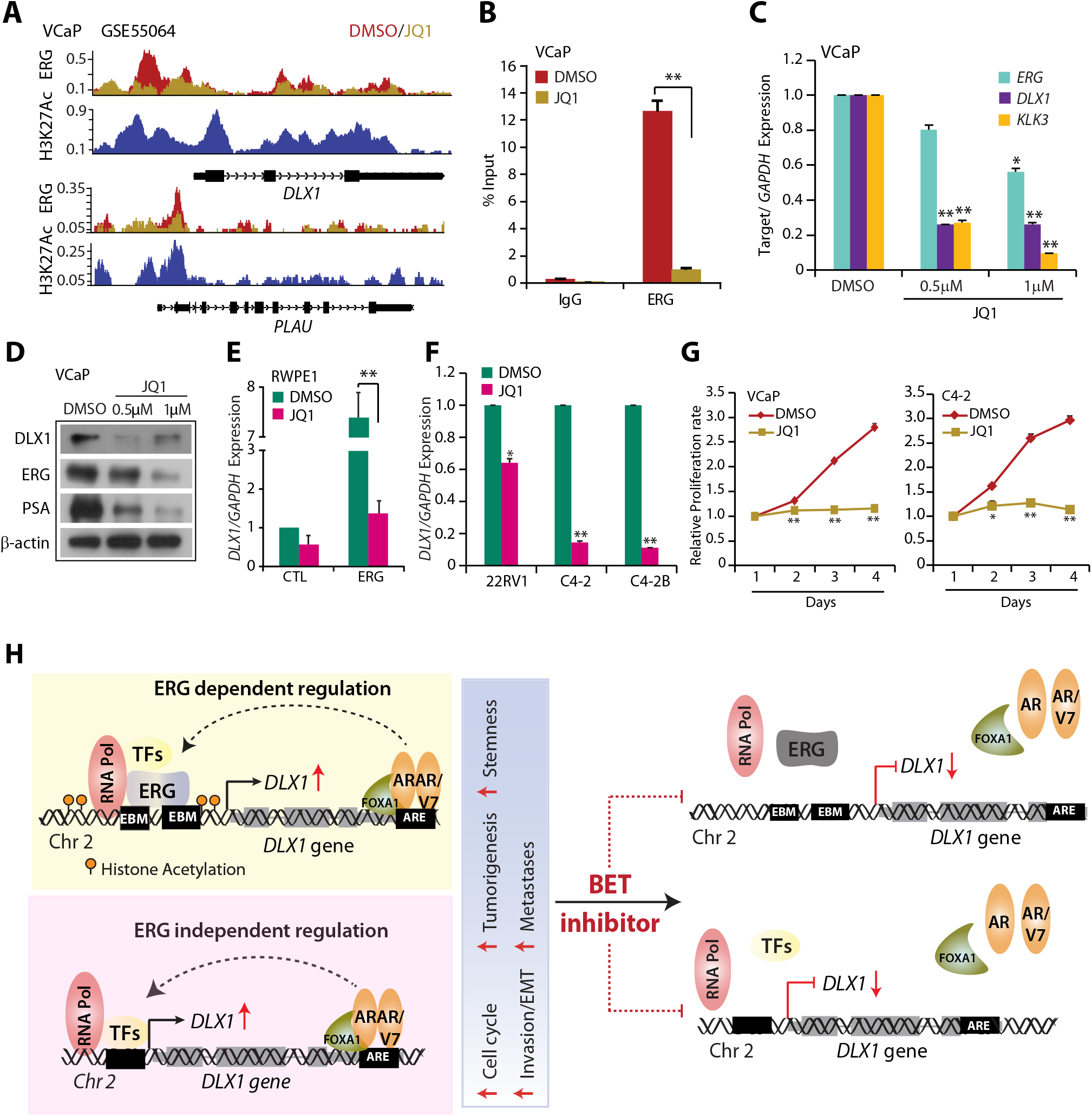
BET domain inhibitor mediates downregulation of DLX1 and its oncogenic properties. **(A)** ChIP-seq data (GSE55064) showing enrichment of ERG on the *DLX1* promoter in VCaP cells treated with JQ1 or vehicle control DMSO for 24 hours. H3K27Ac was used to mark the active promoter region in untreated cells. *PLAU* was used as a positive control for ERG target genes. **(B)** ChIP-qPCR data showing relative enrichment of ERG on the *DLX1* promoter in VCaP cells treated with JQ1 (0.5μM) for 48 hours. **(C)** Quantitative PCR data showing relative expression of target genes in JQ1 treated VCaP cells. *KLK3* used as a positive control for JQ1 treatment. **(D)** Same as in (C) except immunoblot data. β-actin was used as positive control. **(E)** Quantitative PCR showing *DLX1* in RWPE1-ERG and control cells upon JQ1 (0.5μM) treatment. **(F)** Quantitative PCR showing *DLX1* expression in 22RV1, C4-2 and C4-2B cells treated with JQ1 (0.5mM) for 24 hours. **(G)** Cell proliferation assay in VCaP and C4-2 cells upon JQ1 treatment at the indicated time points. **(H)** Schematic showing ERG and AR/AR-V7 dependent transcriptional circuitry regulating DLX1 expression and its oncogenic properties. Also, inhibition of this transcriptional network using BETi results in reduced DLX1 expression and oncogenesis. Data shown from three biologically independent samples. In the panels (B-C), (E), and (G) data represent mean ± SEM. \**P*≤ 0.05 and \*\**P*≤ 0.005 using two-tailed unpaired Student’s t-test.

## Discussion

Detection of *DLX1* along with *HOXC6* in post-DRE urine samples has been introduced as an early diagnostic marker to reduce unnecessary biopsies and to improve the selection of PCa patients at increased risk for high-grade disease (Van Neste et al., 2016). Here, we show that higher expression of DLX1 in PCa patients’ specimens is associated with aggressive disease and poor patient survival. Moreover, using loss-of-function model and cell lines-based assays we demonstrate that DLX1 exhibits pleiotropic cancer effects by engaging into several oncogenic pathways such as proliferative advantage, invasion, EMT and cancer stemness. Importantly, our study demonstrates the positive association between DLX1 and ERG in *TMPRSS2-ERG* positive PCa cases, where patients’ positive for both ERG and DLX1 expression exhibits higher Gleason score and poor survival probability, emphasizing the oncogenic cooperativity which may contribute to disease aggressivity and distant metastases. This is the first report to reveal transcriptional regulation of *DLX1* by ERG oncoprotein in *TMPRSS2-ERG* fusion-positive cases. The TDRD1 (Tudor Domain Containing 1), another established biomarker for PCa has also been shown to be differentially regulated in *ERG* fusion-positive patients, wherein ERG modulates the methylation patterns of the *TDRD1* promoter thereby activating its transcription (Boormans et al., 2013; Kacprzyk et al., 2013). Furthermore, meta-analysis of PCa gene expression data from five independent datasets indicated co-clustering of ERG with DLX1 implicating the hierarchical gene regulatory network of transcription factors, suggesting that ERG might modulate the TFs involved in development (Barfeld, East, Zuber, & Mills, 2014). In corroboration with these independent studies, our findings draw attention to the hierarchical regulatory network between ERG and DLX1, wherein the emergence of *TMPRSS2-ERG* fusion as an early event in primary PCa results in direct transcriptional upregulation of DLX1 via ERG oncoprotein contributing to PCa progression.

AR signaling in CRPC is predominantly dependent on the expression of constitutively active ligand-independent AR-V7 (Dehm, Schmidt, Heemers, Vessella, & Tindall, 2008). Several studies reported AR transcriptional cistrome reprogramming in metastatic CRPC, wherein a cellular context-dependent transcriptional network operates downstream of AR which is distinct from the AR transcriptional program engaged in androgen-dependent stage (Q. Wang et al., 2009). Furthermore, genome-wide AR binding profiles (Q. Wang et al., 2009) demonstrated a highly complex transcriptional circuitry, where AR possibly functions through enhancers distant from the promoters to impart regulation of target genes (Chng et al., 2012; Yu et al., 2010). Although it has been shown that ERG represses AR expression and inhibits AR-mediated transcriptional regulation of canonical genes (Yu et al., 2010), several reports show the cooperativity between these TFs, where AR binds at the enhancer region and forms chromatin loop to interact with ERG at gene promoters thereby regulating the expression of downstream target genes (Cai et al., 2013; Zhang et al., 2019). Recent advancements in understanding the genomic interactions and chromatin looping using high throughput sequencing and conformation capture techniques provided important insight of the high-resolution 3D chromatin landscape of normal and PCa cells (Ramanand et al., 2020). Moreover, the crucial information about mapping of the chromatin interaction of RNA-Pol II, AR and ERG in PCa demonstrates the long- and short-range interactions of these TFs orchestrating genomic expression in PCa (Ramanand et al., 2020; Zhang et al., 2019). In line with this, our findings suggest the formation of active transcriptional complex involving recruitment of AR-FL along with FOXA1 at the putative enhancer region of *DLX1*, which also interacts with ERG occupied *DLX1* promoter in fusion-positive cases, and possibly with other coregulatory TFs in fusion-negative cells. We speculate that this short-range chromatin loop structure between enhancer-promoter region of *DLX1* is further enabled by the RNA-Pol II associated interaction at *DLX1* genomic region, implicating the AR driven transcriptional activation of *DLX1* in an ERG-dependent and -independent manner. Besides, our results also indicate the enrichment of AR-V7 at *DLX1* putative enhancer region and further reveal the role of constitutively activated AR-signaling in regulating *DLX1* expression. Conclusively, these findings demonstrate the role of AR signaling wherein both AR-FL and AR-V7 regulate *DLX1* transcriptional activation in advanced PCa.

It has been known that AR facilitates transcriptional regulation by partnering with transcriptional co-regulators such as histone deacetylases (HDACs), bromodomain-containing proteins (BRDs), histone methyltransferase EZH2 as well as ETS TFs such as ERG (Asangani et al., 2014; Kang, Jänne, & Palvimo, 2004; K. Xu et al., 2012; Zhang et al., 2019). BETi are shown to attenuate AR and ERG-mediated oncogenesis in CRPC cases by disrupting transcriptional activation complex at their target gene loci (Asangani et al., 2014; Blee, Liu, Wang, & Huang, 2016; Chng et al., 2012). Currently, therapeutic targeting of bromodomain proteins have gained clinical importance for the treatment of several malignancies including CRPC, for instance, BETi such as OTX-015, ZEN003694, and GS-5829 are already in clinical trials as single agents or in combination with anti-androgens for CRPC patients (Stathis & Bertoni, 2018). Interestingly, our findings provide convincing evidence that BETi could be used to mitigate the oncogenic effects of DLX1 via disrupting AR and ERG transcriptional circuitries and can be used as a potential therapeutic intervention in treatment of DLX1-positive subtype of PCa patients.

Integrative omics approaches such as Tracing Enhancer Networks using Epigenetic Traits (TENET) have identified DLX1 to be associated with over 100 tumor-specific active enhancers in primary PCa patients, thereby implicating the significance of the extensive DLX1 cistrome that contributes to tumor progression (Rhie et al., 2016). Hence, dissecting downstream regulatory network of DLX1 would be helpful in designing novel therapeutic strategies for the malignancies with higher DLX1 levels. Although *TMPRSS2-ERG* gene fusion is the highly prevalent genetic alteration in PCa, unlike gene fusions involving oncogenic kinases (e.g., RAF-kinase fusions in PCa) (Palanisamy et al., 2010), transcription factors such as ERG are challenging to target. Recent advancements with the development of small molecular inhibitors (Darnell Jr, 2002) and peptidomimetics (X. Wang et al., 2017) show promise to target transcription factors in cancers, thus paving a way to utilize DLX1 as a potential drug target. Collectively, this study provided strong evidence to employ parallel treatment regimen with BETi and AR targeted therapeutics for the clinical management of DLX1-positive subtype, hence opening new treatment avenues for patients with advanced stage disease.

## Materials and Methods

### *In silico* data processing and computational analysis

To study the association of DLX1 with PCa, gene expression data for *DLX1* in TCGA-PRAD dataset was downloaded from UCSC Xena browser (https://xenabrowser.net). The samples were sorted according to the tissue type and plotted for *DLX1* expression (log_2_ (norm_count+1)) in solid normal tissue (n=52) against primary PCa tissue (n=498) using GraphPad prism version 7.0. No cut-off was applied on the dataset. Similar analysis was applied on the dataset retrieved from Gene Expression Omnibus (GEO) database with accession number GSE35988. For GSE80609, analysis was performed as same as for TCGA-PRAD except for FPKM values were plotted. For Kaplan-Meier survival analysis, survival data of patients with primary tumors was considered. Days to first biochemical recurrence and days to last follow-up for TCGA-PRAD patients were the parameters taken into account for the analysis. Samples were divided into two groups based on the expression level of *DLX1* using Cox proportional hazards regression model in R version 3.6.1. 5-year survival probability was calculated using Kaplan-Meier survival analysis (Efron, 1988) by applying survival package (https://cran.r-project.org/web/packages/survival), and log-rank test was used to detect statistical significance. For survival analysis of patients with varying DLX1 expression and its association with other TCGA-defined PCa subtypes (including ERG), data was obtained from UALCAN database (Chandrashekar et al., 2017).

For the heatmap representation, data for *ERG* and *DLX1* was retrieved from TCGA-PRAD using UCSC Xena browser followed by performing hierarchical clustering using heatmap.2 function of gplot package in R version 3.6.1. No cut-off was applied on the dataset. Correlation plot between ERG and DLX1 in TCGA-PRAD and MSKCC dataset was directly retrieved from GEPIA (Tang et al., 2017) and cBioPortal (Gao et al., 2013), respectively.

### Bioinformatics analysis of sequencing data

For RNA-sequencing, raw data (FASTQ format) files for RWPE1, VCaP and 22RV1 were downloaded from GEO with the accession numbers GSE128399 and GSE118206. FastQC sequence quality check was performed on the raw reads followed by sequence mapping using TopHat reader in Galaxy web platform (Afgan et al., 2018) using the default settings. Read alignment results were visualized using Integrative Genomic Browser (IGB) (Freese, Norris, & Loraine, 2016) in reference to the human genome.

ChIP-sequencing data analysis of online available datasets on GEO was performed to determine recruitment of target proteins at *DLX1* promoter region. Galaxy web platform was used to perform ChIP-seq analysis with default settings. Raw single-end reads in FASTQ format were uploaded in the web server using the NCBI SRA accession number for individual samples followed by FASTQC and sequence trimming with FASTQ Trimmer. Next, Sequence Alignment/Map (SAM) files were generated by read alignment to the reference human genome (version as mentioned in the respective study) using Bowtie. Aligned reads were further filtered for the unmapped reads and were converted to the Binary Alignment/Maps (BAM) file format using SAMtools. ChIP-seq peak calling was done using Model-based analysis of ChIP-seq (MACS; *P* < 10^−5^) data with default settings against respective controls. MACS output files were visualized using IGB.

### Cell lines culture conditions and authentication

Prostate cancer cell lines (22RV1, VCaP, LNCaP) and benign prostate epithelial cells (RWPE1) were obtained from the American Type Culture Collection (ATCC) and were cultured as per the ATCC recommended guidelines using growth medium supplemented with 10% fetal bovine serum (FBS) and 0.5% Penicillin-Streptomycin (Gibco Thermo-Fisher), in cell culture incubator supplied with 5% CO_2_ at 37°C.

For authenticating the cell line identity, short tandem repeat (STR) profiling was performed at the Lifecode Technologies Private Limited, Bangalore and DNA Forensics Laboratory, New Delhi. *Mycoplasma* contamination test using PlasmoTest mycoplasma detection kit (InvivoGen) was done routinely for all the cell lines.

### Establishing *DLX1* knockout cell line

CRISPR/Cas9 knockout (KO) kit was purchased from Origene (KN206895) and cell lines were generated following the manufacturer’s protocol. In brief, pCas-guide vector containing guide RNA (gRNA) sequence (specific to *DLX1* and scramble control) along with donor vector containing homologous arms and functional cassette were co-transfected in the host 22RV1 cells. Cells were passaged up to 8 generation post-transfection to eliminate the extrachromosomal donor vector and were then grown under puromycin (Sigma-Aldrich) selection (1μg/ml) pressure to select the positive clones. Single-cell colonies were picked and cultured followed by screening at genomic level. PCR was performed using the primers spanning the deleted genomic region in *DLX1* gene. The *DLX1-KO* cells were further validated for loss of DLX1 at the transcript and protein level using qPCR and immunoblot, respectively (described elsewhere). Primer sequences are provided in Supplementary Table S2.

### Plasmids and Constructs

Briefly, DLX1 cDNA was cloned in pCDH lentiviral vector (Addgene) to establish DLX1 overexpressing cell lines. The pHAGE AR-V7 and control vector were obtained as a generous gift from Dr. Nancy Weigel (Shafi et al., 2015). Successful lentiviral packaging of these constructs was performed to generate stable cell lines (described elsewhere). pGL3-Basic vector, a kind gift from Dr. Amitabha Bandyopadhyay was used to perform promoter reporter assay. For luciferase reporter assays, 1kb promoter region of *DLX1* was cloned in pGL3-Basic vector and site-directed mutagenesis was performed to mutate the ERG binding motifs (EBMs) in *DLX1* promoter. Primer sequences used for mutagenesis are provided in Supplementary Table S2.

### Lentiviral packaging

Lentiviral particles for pCDH vectors were produced using third generation ViraPower Lentiviral Packaging Mix (Invitrogen) according to the manufacturer’s protocol, while for pHAGE vector, second generation pMD.2G and psPAX2 packaging system was procured from Dr. Subba Rao Gangi Setty. Briefly, packaging mix was co-transfected with lentiviral vector in HEK293FT using FuGENE HD Transfection Reagent (Promega). Media containing transfection reagent was replaced with complete growth media after 24 hours. The lentiviral particles were harvested after 48-60 hours, aliquoted and were stored in −80°C freezer.

For establishing stable cell lines, host cells were transduced with viral particles in 6-well dish along with polybrene (hexadimethrine bromide; 8μg/ml) (Sigma-Aldrich) to increase the transduction efficiency. Media was replaced after 24 hours, followed by selection of transduced cells under appropriate antibiotic pressure. To generate DLX1 overexpressing RWPE1 cells, pCDH-DLX1 vector was used for infection and cells were selected in puromycin (0.5μg/ml). LNCaP AR-V7 cells were selected under Geneticin (Gibco, ThermoFisher scientific) at a concentration of 350μg/ml. RWPE1-ERG overexpression and control cells were cultured in Keratinocyte Media with supplements.

### Functional Assays

To perform cell proliferation assay, 1×10^4^ cells were seeded in 12-well cell culture dish and counted using haemocytometer at the indicated time points. Alternatively, Resazurin (Cayman Chemicals) was added to the cells (2×10^3^) plated in 96-well culture dish and fluorescence was measured with emission-excitation at 590-530nm. Relative cell proliferation rate was plotted against the indicated time points.

Foci formation assay was performed by plating 2×10^3^ cells in 6-well culture dish containing the recommended growth media with reduced serum (5% FBS). Cells were incubated at 37°C for three weeks, and media was replaced every third day. Formaldehyde (4% in PBS) was used to fix the cells followed by staining with crystal violet solution (0.05% w/v) and number of foci was counted for quantification.

Cell migration assay was performed using 8μm pore size of Transwell Boyden chamber (Corning). Growth media supplemented with 20% FBS was added to the lower compartment and about 1×10^5^ cells suspended in serum free culture media was added to the Transwell inserts. Cells were incubated at 37°C and after 24 hours migrated cells were fixed in formaldehyde (4% in 1X PBS) and stained using crystal violet (0.5% w/v). Cells adhered on the Transwell filter were detained in 10% v/v acetic acid and migrated cells were quantified by measuring absorbance at 550nm.

For anchorage-independent soft agar assay, 0.6% low melting-point agarose (Sigma-Aldrich) dissolved in RPMI-1640 medium was poured in 6-well culture dishes. After polymerization, another layer of 0.3% soft agar in RPMI-1640 medium containing 22RV1-*DLX1*-KO and 22RV1-SCR control cells (∼1.5×10^4^) was poured on the top of the first layer. Cells were incubated for 20 days at 37°C, and colonies greater than 40μm in size were counted.

### Real Time Quantitative PCR

To identify relative target gene expression, total RNA extracted using TRIzol (Ambion) was converted into cDNA using First Strand cDNA synthesis kit (Genetix) according to the manufacturer’s instructions. Quantitative PCR (qPCR) was performed on the StepOne Real-Time PCR System (Applied Biosystems) using cDNA template, specific primer sets and SYBR Green PCR Master-Mix (Genetix). Primer sequences are provided in the Supplementary Table S2.

### Immunoblotting

Whole cell protein lysates were prepared in RIPA buffer supplemented with Phosphatase Inhibitor Cocktail Set-II (Calbiochem) and protease inhibitor (VWR) followed by protein sample preparation and protein separation on SDS-PAGE. Proteins were then transferred on the PVDF membrane (GE Healthcare) followed by membrane blocking with 5% non-fat dry milk in tris-buffered saline, 0.1% Tween 20 (TBS-T) for 1 hour at room temperature. PVDF membrane was incubated overnight at 4°C with the following primary antibodies at 1:1000 dilution: DLX1 (Thermo, PA5-28899), E-Cadherin (CST, 3195), Vimentin (Abcam, ab92547), phospho-Akt (CST, 13038), total-Akt (CST, 9272), Caspase-3 (CST, 9662), Cleaved PARP (CST, 9541), Bcl-xL (CST, 2764), ERG (Abcam, ab92513), FOXA1 (CST, 58613), 1:2000 diluted AR (CST, 5153), 1:2000 diluted PSA (CST, 5877), and 1:5000 diluted β-actin (Abcam, ab6276). Later, PVDF membranes were washed in washing buffer, 0.1% 1X TBS-T followed by 2 hours incubation at room temperature with secondary anti-mouse or anti-rabbit antibody (1:5000, Jackson ImmunoResearch Laboratories, Cat # 115-035-003 and Cat # 111-035-144, respectively) conjugated with horseradish peroxidase (HRP). Membranes were again washed using 1X TBS-T buffer, and the signals were visualized by enhanced chemiluminescence system (Thermo) according to the manufacturer’s protocol.

### Gene Expression Analysis

Global gene expression profiling and identification of differentially regulated genes was performed as previously described (Bhatia et al., 2018). Briefly, total RNA from 22RV1-*DLX1*-KO and 22RV1-SCR control cells was isolated and subjected to Agilent Whole Human Genome Oligo Microarray profiling (dual color) according to manufacturer’s protocol using Agilent platform (8×60K format). Differentially regulated genes were filtered to only include significantly altered genes with ∼1.6-fold average change (log_2_ fold change greater than 0.6 or less than −0.6, *P*< 0.05). DAVID bioinformatics platform (https://david.ncifcrf.gov) was then used to identify deregulated biological processes. GSEA (Subramanian et al., 2005a) was employed to detect enriched gene signatures upon *DLX1*-KO. Further, network-based analysis of biological pathways was generated using Enrichment Map, a plug-in for Cytoscape network visualization software (Shannon et al., 2003) (http://baderlab.org/Software/EnrichmentMap/).

### Flow Cytometry

For cell cycle distribution, cells were trypsinized and fixed in 70% ethanol followed by staining with propidium iodide (PI) (BioLegend, Cat # 421301) using manufacture’s protocol. For apoptosis, cells were dissociated using StemPro™ Accutase™ (ThermoFisher Scientific) and were washed with cold 1X PBS and resuspended in 1X binding buffer (1×10^6^ cells/ml). Subsequently, 1×10^5^ cells were stained using PE Annexin V Apoptosis Detection Kit I (BD Biosciences, Cat # 559763) following to the manufacture’s protocol. Data was acquired on the Beckman Coulter’s CytoFLEX platform and analyzed using FlowJo software v10.6.1. For apoptotic assay, viable cells were defined by negative stain (lower left quadrant), early apoptotic cells by Annexin V positive and 7-AAD negative (lower right quadrant), dead cells as 7-AAD positive and Annexin V negative (upper left quadrant), and late apoptotic cells as dual positive cells (upper right quadrant).

Aldefluor assay was performed to determine aldehyde dehydrogenase (ALDH) enzymatic activity using Aldefluor kit (Stem Cell Technologies, Catalog #01700) following manufacturer’s guidelines. Briefly, cells were trypsinized and washed with 1X PBS followed by resuspension in 1ml Aldefluor assay buffer. Activated Aldefluor substrate (5μL) was added to the cells and were divided in two conditions with and without ALDH inhibitor, diethylaminobenzaldehyde (DEAB). After 30 minutes of incubation at 37°C, cells were centrifuged and resuspended in 500μL of Aldefluor assay buffer. The ALDH activity was detected in green channel. For data acquisition, Beckman Coulter’s CytoFLEX platform was used and the analysis was performed using FlowJo software v10.6.1.

For stem cell markers, cells were stained with PE/Cy7 anti-human CD44 antibody (Miltenyi Biotec, 130-113-904, 1:50) and CD338-PE (ABCG2-PE, Miltenyi Biotec, 130-105-010, 1:40) followed by 1hour incubation at 4°C. Data acquisition was done using Beckman Coulter’s CytoFLEX platform and analyzed with FlowJo software v10.6.1.

### Mice xenograft study

NOD.CB17-Prkdcscid/J (NOD/SCID) (Jackson Laboratory), five to six weeks old male mice were anesthetized using ketamine and xylazine (50mg/kg and 5mg/kg, respectively) cocktail injected intraperitoneally. 22RV1-*DLX1*-KO and 22RV1-SCR control cells (2×10^6^ for each group) were suspended in 100μl of saline with 20% Matrigel and implanted subcutaneously at the dorsal both flank sides of mice (n=6 for each condition), and tumor burden was measured using digital Vernier caliper on alternate days. Further, tumor volume was calculated using formula (π/6) (L × W^2^), (L=length; W=width).

### Immunohistochemical staining of tumor xenografts

Tumor tissues excised from xenografted mice were fixed in 10% buffered formalin, followed by paraffin embedding and sectioning at 3μm thickness using microtome (Leica). Further, deparaffinization and rehydration of tissue section was performed followed by heat induced antigen-retrieval method using sodium citrate buffer (pH 6.0). Next, endogenous peroxidase activity was quenched using 3% hydrogen peroxide (H_2_O_2_) for 10 minutes, followed by blocking sections in 5% normal goat serum and were probed with primary antibodies against Ki-67 (CST, 9449S), E-Cadherin (CST, 3195), and Vimentin (Abcam, ab92547) at 4°C overnight. Next, after washing sections were incubated with biotinylated secondary antibody at room temperature for 1 hour followed by incubation with ABC (Avidin Biotin complex) (Vector Labs) for 30 minutes. Sections were then processed for detection of HRP activity using DAB (3, 3 -diaminobenzidine) peroxidase (HRP) substrate kit (Vector Labs). Quantification for Ki-67 using 15 random histological fields was performed using imagejs Ki-67 online module (Almeida et al., 2012) and proliferation rate was calculated by normalizing Ki-67 positive nuclei with respect to total nuclei. For quantification of E-cadherin and Vimentin integrated density was calculated using IHC Toolbox in ImageJ software using 15 random histological fields.

### Transcriptome sequencing (RNA-Seq) data analysis

To determine the genes differentially regulated upon siRNA-mediated knockdown of DLX1 in C4-2B cells. We analysed the publicly available RNA-Seq data (GSE78913) using Galaxy (Giardine et al., 2005; Jalili et al., 2020), a web platform available on the public server at https://usegalaxy.org. Raw sequencing FASTQ reads were prefetched and fastq-dumped using SRA-toolkit (http://ncbi.github.io/sra-tools/). Followed by this, reads were assessed for its quality using FASTQC (Andrews, 2010), followed by trimming of data using Trimmomatic (Bolger & Giorgi, 2014). The adapter trimmed reads were next aligned to the human reference genome (hg19) using HISAT2 (Kim, Paggi, Park, Bennett, & Salzberg, 2019) to obtain binary mapped alignment (BAM) files. Next, the transcript abundance among different conditions (siDLX1 treated C4-2B cells and control cells) were analyzed using FeatureCounts (Liao, Smyth, & Shi, 2014). Differential gene expression profiles were evaluated using DESeq2 (Love, Huber, & Anders, 2014) to produce a list of differentially regulated genes with log_2_ fold change (FC) and FDR corrected *P*-values. Genes were annotated using the Entrez gene IDs and those with adjusted *P*-value < 0.05 were selected. Next, differential regulated genes were sorted such that genes with log_2_FC > 0 were considered upregulated while those with log_2_FC < 0 were considered downregulated. We next performed pathway enrichment analysis using DAVID (Sherman & Lempicki, 2009) and GSEA (Subramanian et al., 2005b) to select the pathways deregulated upon *DLX1* knockdown.

### Immunohistochemistry (IHC) and RNA in situ hybridization (RNA-ISH) of PCa patients’ specimens

IHC and RNA-ISH were performed as previously described (Bhatia et al., 2018). Briefly, for IHC, tissue microarray (TMA) slides with PCa tissue samples (n=144) were incubated at 60°C for 2 hours followed by target retrieval in a PT Link instrument (Agilent DAKO, PT200) using EnVision FLEX Target Retrieval Solution, High pH (Agilent DAKO, K800421-2). 1X EnVision FLEX Wash Buffer (Agilent DAKO, K800721-2) was used to wash slides followed by treatment with Peroxidazed 1 (Biocare Medical, PX968M) and Background Punisher (Biocare Medical, BP974L) with a wash for 5 minutes after each step. Slides were incubated with ERG antibody [EPR3864] (Abcam, ab92513) overnight at 4°C. Afterwards, slides were washed and then incubated in Mach2 Doublestain 1 (Biocare Medical, MRCT523L) for 30 minutes at room temperature, and then were rinsed in 1X EnVision Wash Buffer and treated with a Ferangi Blue solution (Biocare Medical, FB813S). Next, slides were rinsed in distilled water followed by treatment with EnVision FLEX Hematoxylin (Agilent DAKO, K800821-2). Slides were dried completely after rinsing in tap water and then dehydrated using xylene. EcoMount (Biocare Medical, EM897L) was added to each slide, which were then mounted with cover slips.

For RNA-ISH, TMA slides incubated at 60°C were de-paraffinized in xylene. Slides were then kept in 100% ethanol twice for 3 minutes each and then air-dried following treatment with H_2_O_2_ for 10 minutes. Further, slides were rinsed and boiled in 1X Target Retrieval for 15 minutes. Slides rinsed in distilled water, were treated with Protease Plus and then incubated with DLX1 probe (Advanced Cell Diagnostics, probe ID: 569601) for 2 hours at 40°C. Slides were washed and treated with Amp 1 for 30 minutes, Amp 2 for 15 minutes, Amp 3 for 30 minutes, and Amp 4 for 15 minutes, all at 40°C in the HybEZ oven with 2 washes in 1X Wash Buffer. Slides were then treated with Amp 5 for 30 minutes and Amp 6 for 15 minutes at room temperature in a humidity chamber. Red color was developed by adding a 1:60 solution of Fast Red B: Fast Red A to each slide and incubating for 10 minutes. Finally, slides were treated with EnVision FLEX Hematoxylin (Agilent DAKO, K800821-2) and mounted using the same protocol as used for IHC slides.

### ERG and *DLX1* staining evaluation

ERG IHC staining was evaluated to define ERG positive and negative status of PCa tissues. *DLX1* expression was identified by scoring the signal intensity of RNA-ISH probe hybridization for the TMA foci and the number of red dots/cells were evaluated to grade *DLX1* expression into four levels ranging from score of 0 to 3 as described previously (Warrick et al., 2014). Next, an association between the expression of ERG and *DLX1* was calculated by applying Chi-Squared contingency test on GraphPad Prism version 7.0.

### Chromatin immunoprecipitation (ChIP)

ChIP was performed as described previously (Tiwari et al., 2020). Briefly, crosslinked cells were lysed using lysis buffer [1% SDS, 10mM EDTA, 50mM Tris-Cl and protease inhibitor (Genetix)] followed by sonication for DNA fragmentation to an average fragment length of ∼500bp using Bioruptor (Diagenode). Sheared chromatin was incubated at 4°C overnight with 4μg of primary or isotype control antibodies. Antibodies against ERG (abcam, ab92513), H3K9Ac (CST, 9733), Rpb1 CTD/RNA PolII (CST, 2629), AR (CST, 5153), control rabbit IgG (Invitrogen, 10500C) and control mouse IgG (Invitrogen, 10400C) were used to perform ChIP assays. Simultaneously, the Protein G coated Dynabeads (Invitrogen) were blocked in presence of BSA (HiMedia) and sheared salmon sperm DNA (Sigma-Aldrich) followed by incubation at 4°C overnight. Blocked beads were incubated for 6-8 hours at 4°C with the lysate containing antibody to make antibody-bead conjugates. Next, the beads conjugated with antibody were washed and immunocomplex was eluted using elution buffer [1% SDS, 100mM NaHCO_3_, Proteinase K (Sigma-Aldrich) and RNase A (500μg/ml each) (Sigma-Aldrich)]. DNA was isolated using phenol-chloroform-isoamyl alcohol extraction method.

For Re-ChIP, first ChIP was performed using ERG antibody eluted in 10mM DTT in Tris-EDTA buffer at 37°C for 30 minutes, subsequently DTT-elute was further diluted 50 times and second round of ChIP was performed following the similar protocol using antibody against AR. The ChIP-qPCR was performed using primers provided in Supplementary Table S2. RNA-Pol II ChIA-PET data (GSM3423997) was downloaded from GEO and visualized using integrated genome viewer (IGV) (Robinson et al., 2011).

### Luciferase Assay

RWPE1-CTL and RWPE1-ERG cells plated in 24-well culture dish were transfected with pGL3-DLX1-P wildtype (250ng) and pRL-null vector (2.5ng) using FuGENE HD Transfection Reagent. After 48 hours of incubation at 37°C, cells were harvested using the lysis buffer provided with Dual-Glo Luciferase assay kit (Promega). GloMax® 96 Microplate Luminometer (Promega) was used to measure the Firefly and Renilla luciferase activity according to the manufacturer’s protocol. Renilla luciferase activity was used as a normalization control. Same protocol was followed to measure the luciferase promoter reporter activity for pGL3-DLX1-P mutant construct.

### Androgen stimulation and compound treatment

For androgen stimulation, cells were starved for 72 hours in phenol-red free media supplemented with 5% charcoal stripped serum (CSS) (Gibco) followed by stimulation with 10nM R1881 (Sigma-Aldrich) at the indicated time points. For anti-androgen treatment, VCaP cells were cultured in phenol-red free DMEM media supplemented with GlutaMAX (Gibco) and 5% charcoal stripped serum (CSS) followed by pre-treatment with antiandrogen for 6 hours. Next, cells were stimulated with 1nM R1881 in presence of anti-androgen for 48 hours. For LNCaP cells anti-androgen treatment was performed similar to VCaP cells, except LNCaP cells were cultured in RPMI-1640 phenol-red free medium (Gibco) supplemented with 5% CSS. For ChIP-qPCR experiments, VCaP and 22RV1 cells were stimulated with 10nM R1881 for 16 hours. For VCaP anti-androgen ChIP experiment, cells were pre-treated with Enzalutamide for 6 hours followed by treatment with R1881 in presence of anti-androgen for 16 hours. For JQ1 treatment, cells were grown in complete growth media followed by JQ1 treatment using indicated concentration for 48 hours.

### Study Approval

In vivo mice procedures were approved by the Committee for the Purpose of Control and Supervision of Experiments on Animals (CPCSEA) and abide to all regulatory standards of the Institutional Animal Ethics Committee of the Indian Institute of Technology Kanpur. TMAs comprising prostate cancer specimens (n=144) were obtained from Department of Pathology, Henry Ford Health System (Detroit, MI), after getting written informed consent and Institutional Review Board approval. All patients’ specimens used in this study were collected in accordance with the Declaration of Helsinki. The patients were not administered with any hormone therapy, except for three patients.

### Statistical Analysis

For statistical analysis, unpaired, two-tailed Student’s *t-*test was used for independent samples or otherwise mentioned. For *P*-value less than 0.05, the differences between the groups were considered significant. Significance was indicated as follows: \**P*<0.05, \*\**P*<0.001. Error bars represent standard error of the mean (SEM) obtained from experiments performed at least three independent times.

## Acknowledgments

This work is supported by the Science and Engineering Research Board (SERB), Government of India (EMR/2016/005273 to B.A.). We also thank research funding from the Wellcome Trust/ DBT India Alliance (IA/I(S)/12/2/500635 to B.A.) and Department of Biotechnology, Government of India (BT/PR8675/GET/119/1/2015 to BA). N.P acknowledges financial support from CMDRP (W81XWH-16-1-0544). We are thankful to Nancy Weigel for providing pHAGE AR-V7 construct, Paul Rennie for the C4-2 shAR cells and Amitabha Bandyopadhyay for sharing pGL3 basic vector. We also thank Jonaki Sen and Pradip Sinha for extending the use of their microscopy facility and Nishat Manzar for critically reading the manuscript.

## Author Contributions

S.G. and B.A. conceptualized the study and designed the experiments. S.G. performed *in vitro* cell line-based studies. S.G. and V.B. performed the gene expression studies, bioinformatics analysis and ChIP assays. S.G. and V.B. executed the *in vivo* mice xenograft studies. S.G. and B.A. performed statistical analysis and interpreted the data. S.C., N.G., and N.P. performed immunohistochemistry and RNA *in situ* staining on the PCa tissue microarrays. M.A. helped in designing experiments related to androgen signaling mediated regulation. S.G. and B.A. wrote the manuscript. B.A. directed the overall project.

