## Supplementary Figures for "Transcriptional network involving ERG and AR orchestrates Distal-Less Homeobox 1 mediated prostate cancer progression"

### Figure Supplement

#### Figure S1

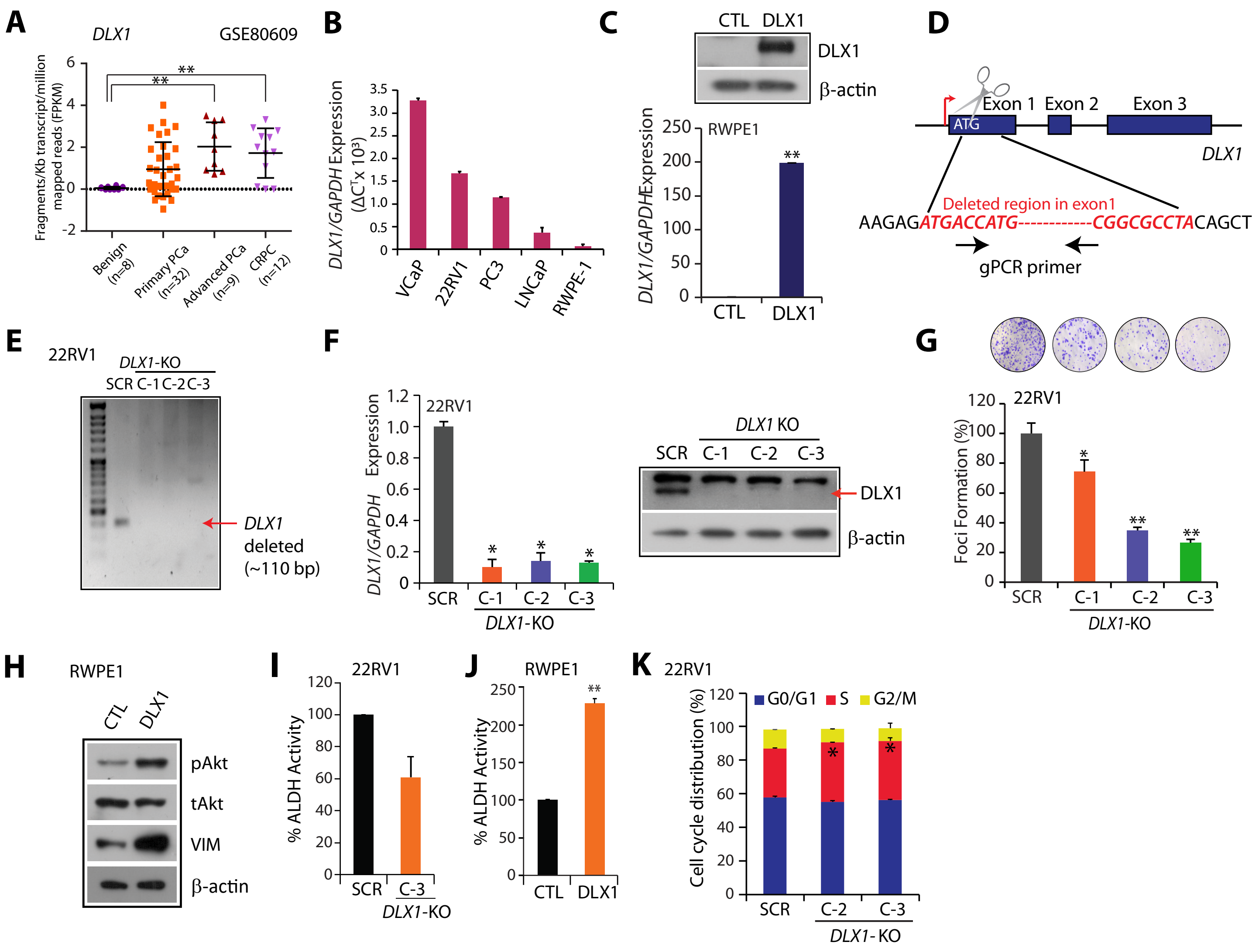


**Figure S1. Characterization of DLX1 overexpression and knockout cells.**
**(A)** Dot-plot of RNA-seq analysis of dataset GSE80609 for *DLX1* in PCa stages, data represents mean ± SD. One-way ANOVA with post hoc Dunnett's multiple comparisons test was applied. **(B)** Quantitative PCR data showing *DLX1* expression in PCa cell line panel. **(C)** Q-PCR and immunoblot showing relative expression of DLX1 in RWPE1-DLX1 overexpressing cells. **(D)** Schematic representation of DLX1 genomic deletion to generate 22RV1-DLX1-KO cells. **(E)** Agarose gel depicting deleted region of *DLX1* in KO clones and control. **(F)** Q-PCR and immunoblot analysis showing DLX1 expression in 22RV1-DLX1-KO cells. **(G)** Foci formation assay using same cells as in (F). Inset shows representative image of foci. **(H)** Immunoblot showing expression of phospho (p) and total (t) Akt and Vimentin in RWPE1-DLX1 cells. β-actin is used as a loading control. **(I)** Bar graph showing percent ALDH activity of the aldeflour flow cytometery assay in *DLX1*-KO and control cells. **(J)** Same as (I) except DLX1 overexpressing RWPE1 cells were used. **(K)** Bar graph representing cell cycle phase distribution in 22RV1-*DLX1*-KO and control cells.
Biologicaly independent samples n=3 were used for the experiments. Data represent mean ± SEM. *$P \leq 0.05$ and **$P \leq 0.005$ using two-tailed unpaired Student's *t*-test, unless specified.

#### Figure S2

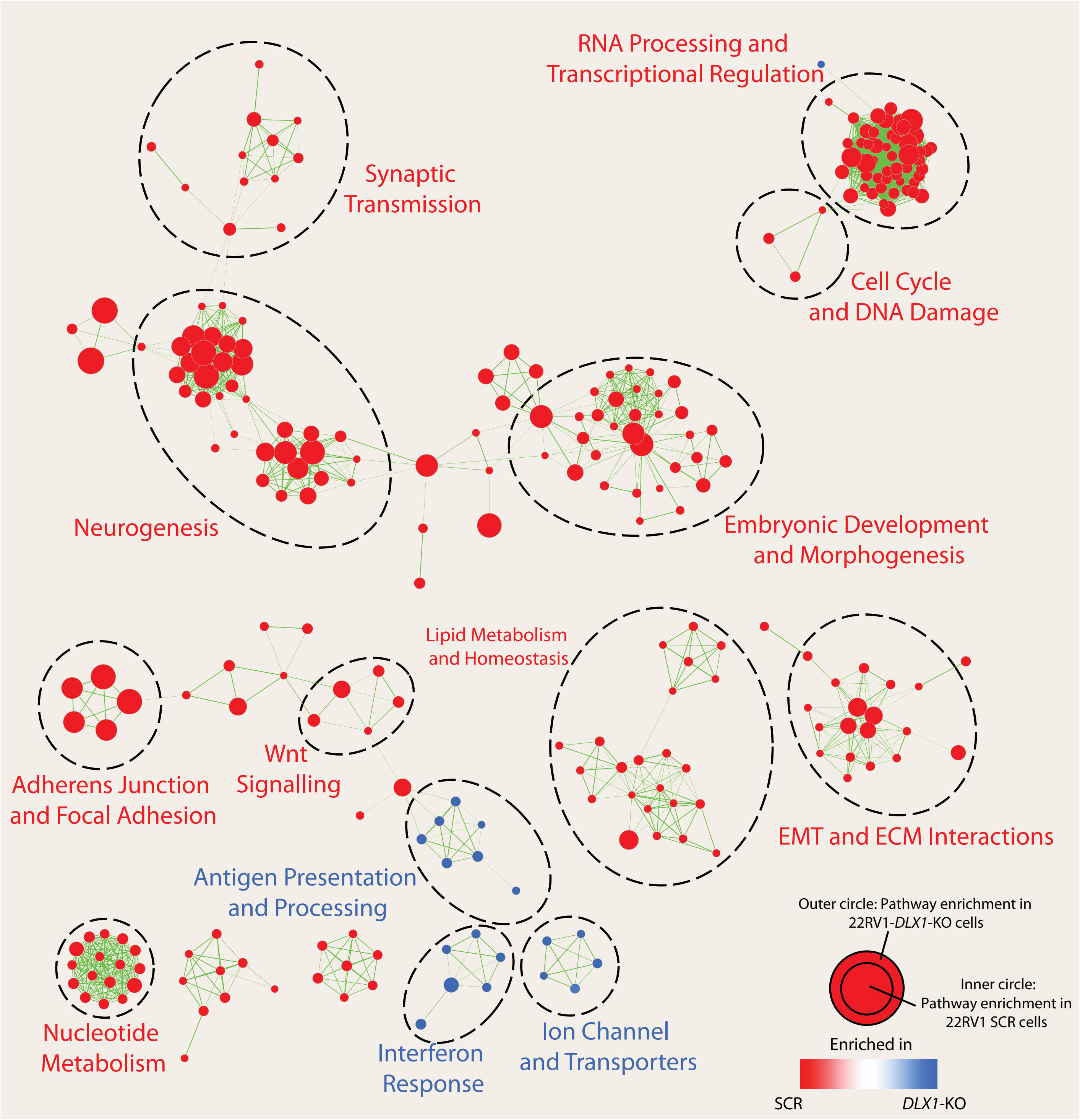


**Figure S2. Enrichment map showing overlapping biological networks.**
GO term enrichment map for the gene expression profiles obtained from microarray analysis of 22RV1-*DLX1*-KO and control cells. Each node here represents a GO term and size of the node correspond to the number of genes in the particular GO term. Red colour indicates the pathways enriched in control group while blue colour shows pathways enriched in *DLX1* knockout cells. Intensity of the colour represents the enrichment significance. Each cluster is manually labelled to depict the pathway enrichment.

### Figure S3

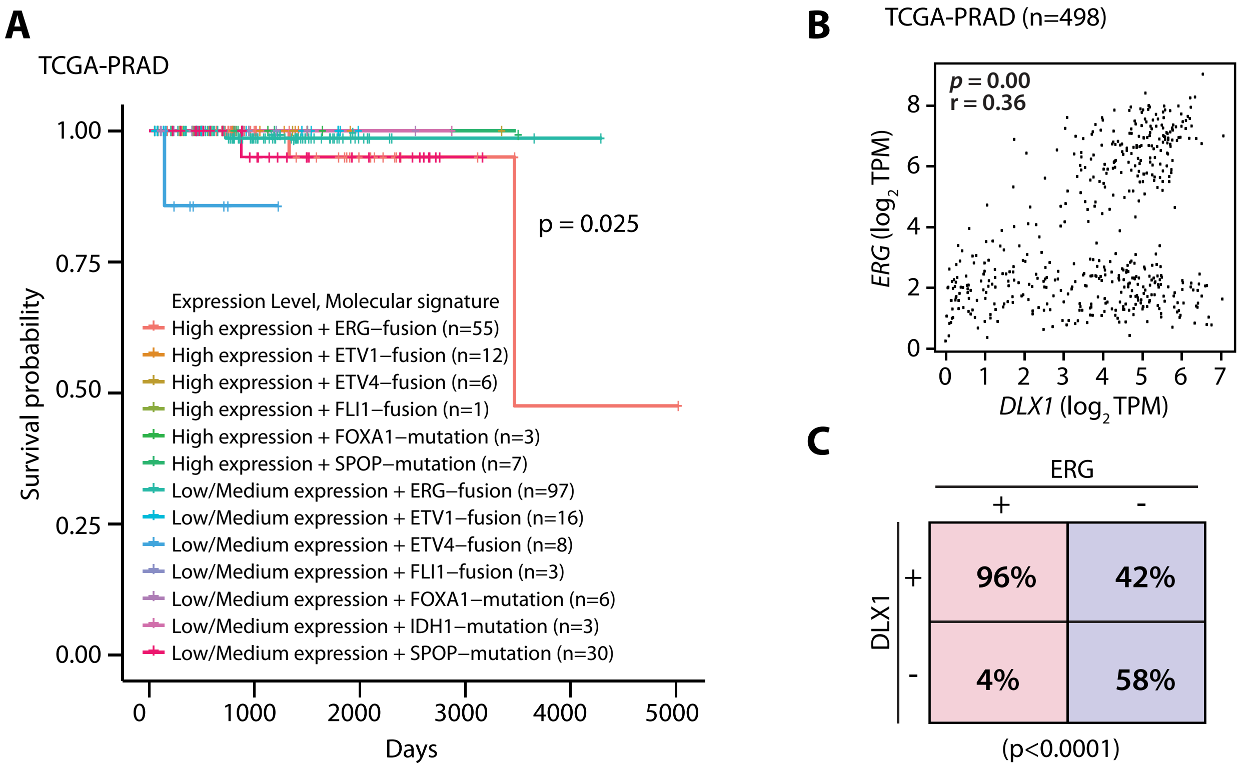


**Figure S3. DLX1 and ERG shows significant positive correlation in PCa patients.**
**(A)** Kaplan Meier plot depicting survival probability of *DLX1* expression level in association of molecular subtypes defined by TCGA in primary prostate cancer patients. **(B)** Correlation plot between *DLX1* and *ERG* transcript per million readcount (TPM) in TCGA-PRAD dataset generated using GEPIA (Gene Expression Profiling Interactive Analysis). **(C)** Contingency table depicting status of *DLX1* and ERG. Red panel shows status of *DLX1* patients in ERG positive cases (left) and blue panel shows status of *DLX1* in ERG negative cases (right). P-value for Fisher's exact test is indicated.

### Figure S4

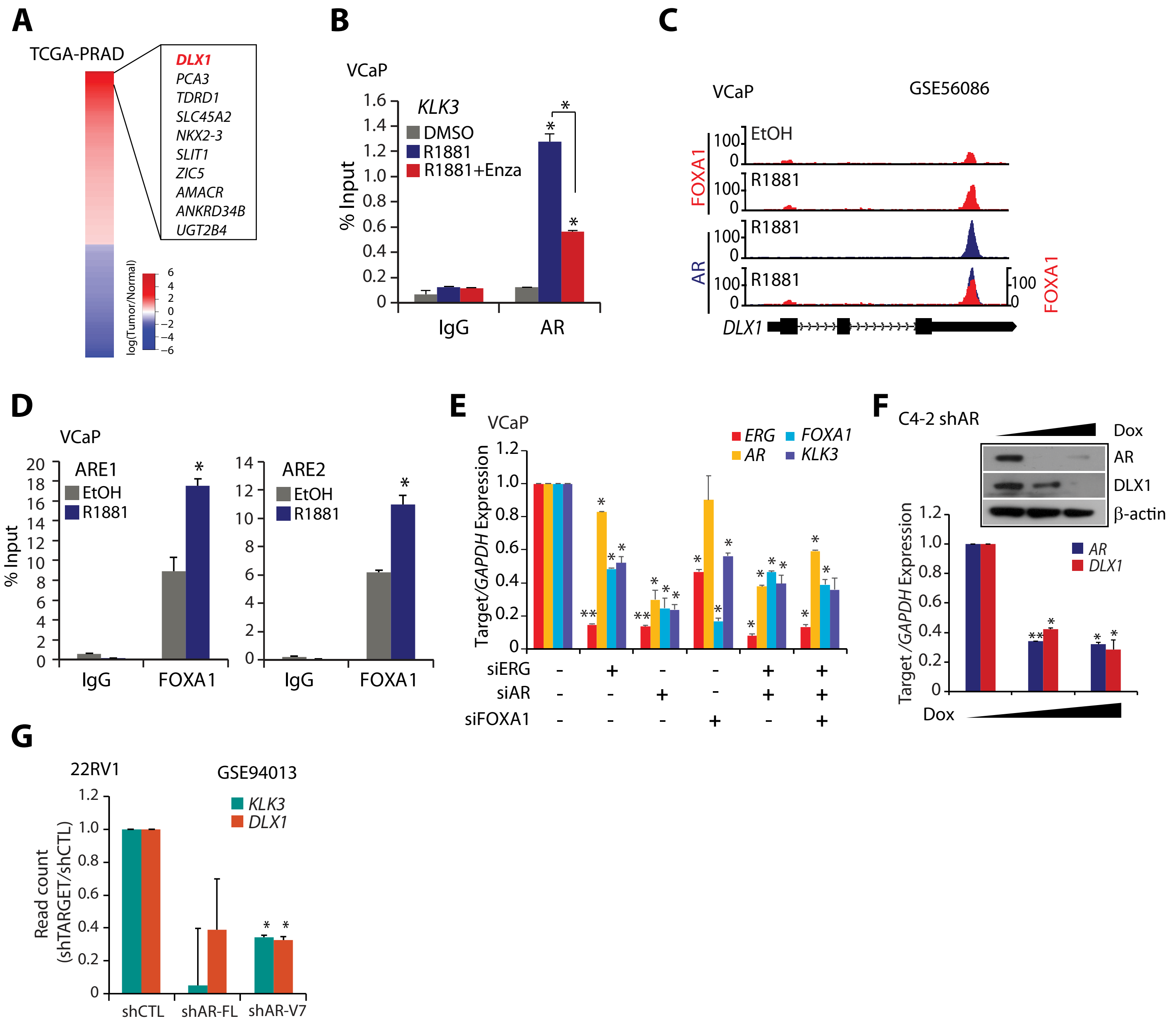


**Figure S4. AR gets recruited at *DLX1* putative enhancer in PCa cells.**

**(A)** Heatmap showing top 10 differentially upregulated genes in TCGA-PRAD dataset with tissue specific AR binding site in 50kb nearby region. **(B)** ChIP-qPCR data showing AR recruitment at *KLK3* promoter in VCaP cells treated with vehicle control, R1881 or combination of R1881 and Enza. **(C)** ChIP-seq data depicting recruitment of AR and FOXA1 at the same genomic loci of *DLX1* gene in R1881 stimulated VCaP cells. **(D)** ChIP-qPCR data depicting FOXA1 recruitment at the DLX1 putative enhancer in R1881 (10nM) stimulated VCaP. **(E)** Quantitative PCR depicting relative expression of target genes in VCaP cells transfected with siRNA against AR/ERG and/or FOXA1. **(F)** Q-PCR and immunoblot for AR and DLX1 in doxycycline (dox) inducible shAR C4-2 cells treated with dox for 48 hours. **(G)** Bar plot showing relative read count of *KLK3* and *DLX1* obtained from RNA-seq data for *AR-FL* and *AR-V7* knockdown in 22RV1 cells downloaded from GEO (GSE94013).

Biologically independent samples n=3 were used for the experiments. Data represent mean±SEM. *$P \le$ 0.05 and **$P \le$ 0.005 using two-tailed unpaired Student's *t*-test.
