## Supplementary Table S2 for "Transcriptional network involving ERG and AR orchestrates Distal-Less Homeobox 1 mediated prostate cancer progression"

**Supplementary Table S2:** List of Primers

| **Quantitative PCR (qPCR) primers** | | | |
|---|---|---|---|
| S.No. | Gene | Primer Name | |
| 1 | *DLX1* | qDLX1_FP | GCG GCC TCT TTG GGA CTC ACA C |
| | | qDLX1_RP | GGC CAA CGC ACT ACC CTC CAG A |
| 2 | *CDH1* | qCDH1_FP | CTT CTG CTG ATC CTG TCT GAT G |
| | | qCDH1_RP | TGC TGT GAA GGG AGA TGT ATT G |
| 3 | *VIMENTIN* | qVIM_FP | GAT TCA CTC CCT CTG GTT GAT AC |
| | | qVIM_RP | GTC ATC GTG ATG CTG AGA AGT |
| 4 | *SNAIL* | qSNAIL_FP | CCT TCG TCC TTC TCC TCT ACT T |
| | | qSNAIL_RP | TTC GAG CCT GGA GAT CCT T |
| 5 | *POU5F1/Oct-4* | qOCT_FP | GGA GGA AGC TGA CAA TGA AA |
| | | qOCT_RP | GGC CTG CAC GAG GTT TT |
| 6 | *ABCG2* | qABCG2_FP | GTA AAG CAG GGC ATC GAT CT |
| | | qABCG2_RP | CAG GTA GGC AAT TGT GAG GAA |
| 7 | *C-KIT* | qKIT_FP | CAA GGC TTC TCC AAT TCT GC |
| | | qKIT_RP | TGC AGT GGT CCA CAG AAG AG |
| 8 | *SOX2* | qSOX2_FP | CAT GGG TTC GGT GGT CAA G |
| | | qSOX2_RP | TGA TCA TGT CCC GGA GGT |
| 9 | *ERG* | qERG_FP | CGC AGA GTT ATC GTG CCA GCA GAT |
| | | qERG_RP | CCA TAT TCT TTC ACC GCC CAC TCC |
| 10 | *KLK3* | qPSA_FP | GTC TGC GGC GGT GTT CTG |
| | | qPSA_RP | TGC CGA CCC AGC AAG ATC |
| 11 | *AR* | qAR_FP | AAT CCC ACA TCC TGC TCA AG |
| | | qAR_RP | GAG TCC AGG AGC TTG GTG AG |
| 12 | *FOXA1* | qFOXA1_FP | ATA CTC GCC TTA CGG CTC TA |
| | | qFOXA1_RP | GTT TAG GAC GGG TCT GGA ATA C |
| 13 | *GAPDH* | qGAPDH_FP | TGC ACC ACC AAC TGC TTA GC |
| | | qGAPDH_RP | GGC ATG GAC TGT GGT CAT GAG |
| **Chromatin Immunoprecipitation (ChIP-qPCR) primers** | | | |
| 1 | *KLK3* | KLK3_FP | CCT AGA TGA AGT CTC CAT GAG CTA CA |
| | | KLK3_RP | GGG AGG GAG AGC TAG CAC TTG |
| 2 | *DLX1* | EBM1_FP | AGC TTT GAA CCG AGT TTG GG |
| | | EBM1_RP | TTC TCT CCT CTG CTT CCC TTT |
| 3 | *DLX1* | EBM2_FP | CAG CCC ATT GTG CTT CCT G |
| | | EBM2_RP | GGT CCG CTG TCT TGC ATA ATC |
| 4 | *DLX1* | ARE1_FP | CGC CTG GAC AAG AAG GAA A |
| | | ARE1_RP | CCA CTG GAA TTA GAG TGC TTG T |
| 5 | *DLX1* | ARE2_FP | GAG AAA TGG ACT TCG CCT GA |
| | | ARE2_FP | CCC GTG CGC TTA AAG TAA AC |
| 6 | *CUTL2* | CUTL2_FP | AAA CAT GTC TCC CCA TGG AA |
| | | CUTL2_RP | GGT ACA TCC TGC ACC AGA CC |
| **Genomic PCR primers** | | | |
| 1 | *DLX1* | gDLX1del_FP | ATG ACC ATG ACC ACC ATG CCA |
| | | gDLX1del_RP | AAT GGC CCG CCG AGT GTA AA |
| **Mutagenesis Primers** | | | |
| 1 | *DLX1* | MT1_FP | CCA TTG TGC AAC CTG CCC GG |

| | | | |
|---|---|---|---|
| | | MT1_RP | CCG GGC AGG TTG CAC AAT GG |
| 2 | *DLX1* | MT2_FP | TTG GGT TCC AGC CTG TCC TGA |
| | | MT2_RP | TCA GGA CAG GCT GGA ACC CAA |

**Data file S1**: Deregulated biological processes in 22RV1-*DLX1*-KO cells predicted by DAVID analysis.
